# High Throughput Computational Mouse Genetic Analysis

**DOI:** 10.1101/2020.09.01.278465

**Authors:** Ahmed Arslan, Yuan Guan, Zhuoqing Fang, Xinyu Chen, Robin Donaldson, Wan Zhu, Madeline Ford, Manhong Wu, Ming Zheng, David L. Dill, Gary Peltz

## Abstract

**Background:** Genetic factors affecting multiple biomedical traits in mice have been identified when GWAS data that measured responses in panels of inbred mouse strains was analyzed using haplotype-based computational genetic mapping (HBCGM). Although this method was previously used to analyze one dataset at a time; but now, a vast amount of mouse phenotypic data is now publicly available, which could lead to many more genetic discoveries.

**Results:** HBCGM and a whole genome SNP map covering 53 inbred strains was used to analyze 8462 publicly available datasets of biomedical responses (1.52M individual datapoints) measured in panels of inbred mouse strains. As proof of concept, causative genetic factors affecting susceptibility for eye, metabolic and infectious diseases were identified when structured automated methods were used to analyze the output. One analysis identified a novel genetic effector mechanism; allelic differences within the mitochondrial targeting sequence affected the subcellular localization of a protein. We also found allelic differences within the mitochondrial targeting sequences of many murine and human proteins, and these could affect a wide range of biomedical phenotypes.

**Implications:** These initial results indicate that genetic factors affecting biomedical responses could be identified through analysis of very large datasets, and they provide an early indication of how this type of ‘*augmented intelligence*’ can facilitate genetic discovery.

## Introduction

Mouse is the premier model organism for biomedical discovery, and mice were used for the discovery or development of many therapies that are now in clinical use. However, similar to the difficulties encountered in analyzing human GWAS results [1], it has been difficult to identify the genetic factors underlying biomedical trait response differences in GWAS using inbred mouse strains. Just as human subpopulations are descended from ancestral founders; inbred laboratory strains are derived from an estimated four ancestral founders [2, 3]. Because of their ancestral relatedness, GWAS results will identify a true causative variant along with multiple other false positive associations, which are caused by genomic regions with a correlated genetic pattern that is due to common inheritance (a property referred to as ‘population structure’). Statistical methods have been developed to reduce the false discovery rate by correcting for the population structure that exists in humans [4, 5], plants [6], and mice [7]. While these correction methods have utility for analysis of human GWAS results, we have shown that they are less useful for analyzing murine GWAS results; and moreover, their use could also increase the chance that a true causative genetic factor will be discarded [8]. In brief, even though multiple genomic regions have a shared ancestral inheritance, one may be responsible for a phenotypic difference.

Haplotype-based computational genetic mapping (**HBCGM**) [9] is a method for analyzing mouse GWAS data, which has identified genetic factors underlying 22 biomedical traits in mice [9-31]. One finding generated a new treatment for preventing narcotic drug withdrawal [23] that is now being tested in a multi-center clinical trial [32]. In an HBCGM experiment, a property of interest is measured in available mouse strains whose genomes have been sequenced; and genetic factors are computationally predicted by identifying genomic regions (haplotype blocks) where the pattern of within-block genetic variation correlates with the distribution of phenotypic responses among the strains [10, 33, 34]. A next-generation version of HBCGM with a 30,000-fold improvement in computational efficiency was developed, and whole genome sequence data for 26 strains was analyzed to produce a whole genome map with 16M SNPs [15]. HBCGM was previously used to analyze the response data for one trait at a time. But now, a vastly increased amount of phenotypic data for inbred mouse strains has become available. The Mouse Phenome Database (**MPD**) [35] has 8462 phenotypic datasets (1.52M individual datapoints) that measure experimentally-induced responses in panels of inbred mouse strains. They have been shown to be useful for genetic discovery since a genetic susceptibility factor for haloperidol-induced toxicity was identified by analysis of one MPD dataset [15]. Many more genetic discoveries could be made if all 8462 of the MPD datasets could be analyzed. However, HBCGM analyses generate many false positive associations that appear along with the causative genomic region for the trait response difference in the list of correlated genes. This creates a significant hurdle that limits our ability to genetically analyze the information contained within a large database like the MPD. For example, if 50 correlated genomic regions were identified for each of the 8462 MPD datasets (i.e., 415K possible genetic leads), the output could not be carefully examined by a team of dedicated individuals (or even by a University’s entire staff of scientists). This problem is compounded by the fact that the HBCGM program maintains a relatively low threshold for identification of correlated genetic regions to avoid false negatives. To overcome this problem, selection methods are used to identify the true causative factor from among the multiple correlated regions. Causative genetic candidates were selected from among the many genes with correlated allelic patterns by applying orthogonal criteria [10, 36], which include gene expression, metabolomic [22], or curated biologic data [37], or by examining candidates within previously identified genomic regions [24, 25]. This approach evaluates genetic candidates using multiple criteria; this can provide superior results to that of a typical GWAS that only uses a single highly stringent criterion to identify candidates. In order to more efficiently identify likely candidate genes, the logical paths that were used to select the previously identified genetic factors were used to develop structured computational methods for analyzing HBCGM output. We demonstrate the utility of this approach by identifying several murine genetic factors that were known to affect important biomedical phenotypes. This approach is then used to identify a novel genetic effector mechanism that alters the expression and subcellular localization of the encoded protein. We also find that this novel genetic effector mechanism could affect many mouse and human genes.

## Results

### Generation and characterization of the SNP database

Whole genome sequencing data was used to generate a database with 21.3M SNPs with alleles covering 53 inbred mouse strains (**Table S1**). The fold genome coverage averages 41.6x per strain (range 19x to 168x) based upon a 3 Gb genome); which is similar to that of other recent studies [38, 39]. The high fold-genome coverage and the use of stringent tiered methods used for variant calling [15] ensured that variants were identified with high confidence. However, to evaluate the quality of the identified SNP alleles, we compared our variant calling method (BCFtools) with the results obtained using the GATK HaplotypeCaller [40]. To do this, the alleles identified by the two methods were compared with those in the NCBI dbSNP (build 142), which was downloaded from mouse genome informatics (MGI) site (http://www.informatics.jax.org/snp). We evaluated SNPs identified on chromosome 1 by the two different methods for 32 of the inbred strains. the BCFtools variant calling results agreed with those in dbSNP ∼90% of the time (**Fig. S1A**). The rates of correct calls for GATK and BCFtools were stable across the 32 inbred strains and were independent of the depth of sequence coverage (**Fig. S1B**). The rates of incorrect and missed calls generated using the BCFtools or GATK pipeline were low (<8%), while GATK has a higher rate of missed calls than BCFtools. The average concordance rate for all SNP allele calls across the 32 strains was above 97%. This assessment of the BCFtools pipeline performance provides confidence in the SNP database that is used for HBCGM analyses.

### Bulk analysis of MPD datasets

Each of the 8462 MPD datasets is categorized according to the type of biomedical trait response measured [35]. As examples, 472 datasets relate to body weight or body fat composition; 255 measure an immune system response; 96 relate to drug metabolism; and 233 datasets measure a neurologic response. In some cases, multiple datasets measure the same response at different times after a treatment (i.e. haloperidol toxicity [15]). Irrespective of their grouping, we identified 2138 MPD datasets that measured a response in 12 or more strains and had inter-strain differences that were likely to be heritable (ANOVA p value <1×10^−5^). We found that the selected 2138 MPD datasets had an average of 24 inbred strains that were in our database. After these datasets were bulk analyzed by HBCGM, 1173 datasets had at least one gene with a genetic association p value <1×10^−5^. Given this large number of datasets, we analyzed a few to determine if causal genetic factors could be identified. We present the results from several of these analyses; and show how the use of additional structured computational analysis methods enabled causative genetic factors to be identified.

### Using structured computational methods

One MPD data set (MPD:1501) characterized the susceptibility of bone marrow macrophages obtained from female mice of 23 inbred strains to the *Bacillus anthracis* lethal factor [41]. This toxin killed macrophages obtained from twelve strains, while macrophages from 11 other strains were resistant to the toxin. HBCGM analysis indicated the allelic pattern that was most highly correlated with susceptibility (ANOVA p value = 4.1 ×10^−10^) to this toxin was within the pyrin domain of the *NACHT, LRR and PYD domains containing protein 1b* (*Nlrp1b*) (**Fig 1**). The susceptible and resistant strains had distinct *Nlrp1b* haplotypes. Of note, all of the most highly correlated genes were located in the same region of mouse chromosome 11; and the second most highly correlated gene was an adjacent functional homologue (*Nlrp1a*) of Nalp1b. Nlrp1b has been shown to play a crucial role in the formation of a subset of inflammasomes, which is an important part of the response to pathogenic infections [42, 43]; and allelic variation in *Nlrp1b* was previously shown to regulate macrophage [41] and neutrophil-dependent responses [44] to anthrax infection among inbred strains.

**Figure 1.**
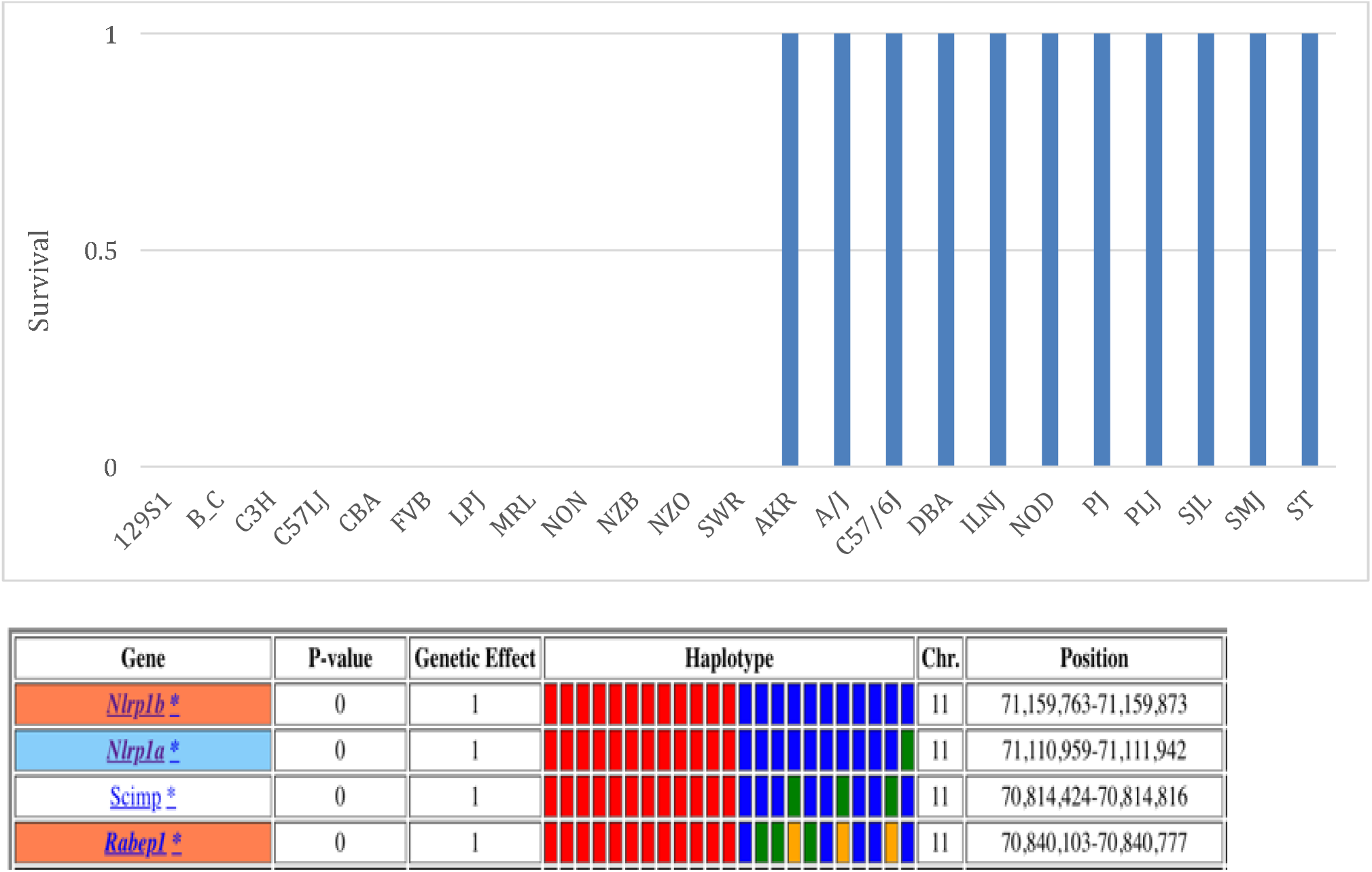
**Top**: The susceptibility of inbred strains to the lethal factor produced by *Bacillus anthracis*. Macrophages isolated from each indicated strain were incubated lethal toxin, and their survival was measured as described [41]. Strains with a “blue bar” (100% survival) are resistant to the toxin, while strains without a bar (100% lethality) were susceptible. **Bottom:** HBCGM analysis identifies the genes whose allelic patterns were most highly correlated with toxin susceptibility. The four co-linear genes with haplotype blocks that correlated with the phenotypic response pattern are indicated by their symbol; and an orange, white, or blue background indicates whether a SNP causes or does not cause an amino acid change, or if it affects a splice site, respectively. The haplotypic pattern is shown as colored rectangles that are arranged in the same order as the input data (shown in the top graph). Strains with the same colored rectangle have the same haplotype within the haplotype block within the indicated gene. The p-values and genetic effect size were calculated as previously described [33].

The anthrax toxin response was binary (death or survival), it was measured in a large number of strains, and the different types of response was evenly distributed among the 23 strains analyzed. However, many MPD datasets examined a smaller number of strains, and the different response types of were unevenly distributed among the strains. Because of this, HBCGM analysis of most MPD datasets identified a much larger number of genes with haplotypic patterns that were highly correlated with the phenotypic response pattern; and structured computational methods for filtering the output were needed to identify the true causative genetic factor. For the cases described here, automated methods were used to identify correlated genes that: (i) were expressed within the target organ for the trait; (ii) contained a codon-changing SNP (cSNP); and (iii) a search of the biomedical literature indicated that the gene was related to the phenotype. For example, one MPD data set (MPD: 26721) examined the retinas of 29 strains: 21 strains had normal retinas, and 8 strains had retinal degeneration (**Fig. 2**). Thirty-three genes had haplotype blocks with a perfect genotype-phenotype correlation (ANOVA p value = 0): all strains with normal retinas had one haplotype, while those with retinal degeneration shared a second haplotype. The computational filtering process rapidly reduced the number of candidates to four genes, which were all located within a single chromosomal region. Of these four genes, only one gene had a SNP allele with a stop codon, and the published literature analysis indicated that it was associated with vision *Phosphodiesterase 6b* (*Pde6b*) encodes a phosphohydrolase that plays key role in transducing light mediated retinal signals [45]. A SNP with a stop codon (*Tyr347X*) was located within a protein domain (GAF) that occurs in cGMP-regulated phosphodiesterases that are responsible for high affinity, non-catalytic binding of two cGMP molecules/holoenzyme. All eight strains with retinal degeneration had the stop codon, while the 29 strains with normal retinas had the *Tyr347* allele (Fig. 2). Although we did not know it when this analysis was performed, a form of retinal degeneration that occurred in some inbred mouse strains was first described 93 years ago [46]; and the causative mutation (*retinal degeneration 1, rd1*) was subsequently localized to this *Pde6b* SNP [47]. Although a previously known genetic factor was identified in this example, this result demonstrates these structured computational analysis methods have utility for genetic discovery.

**Figure 2.**
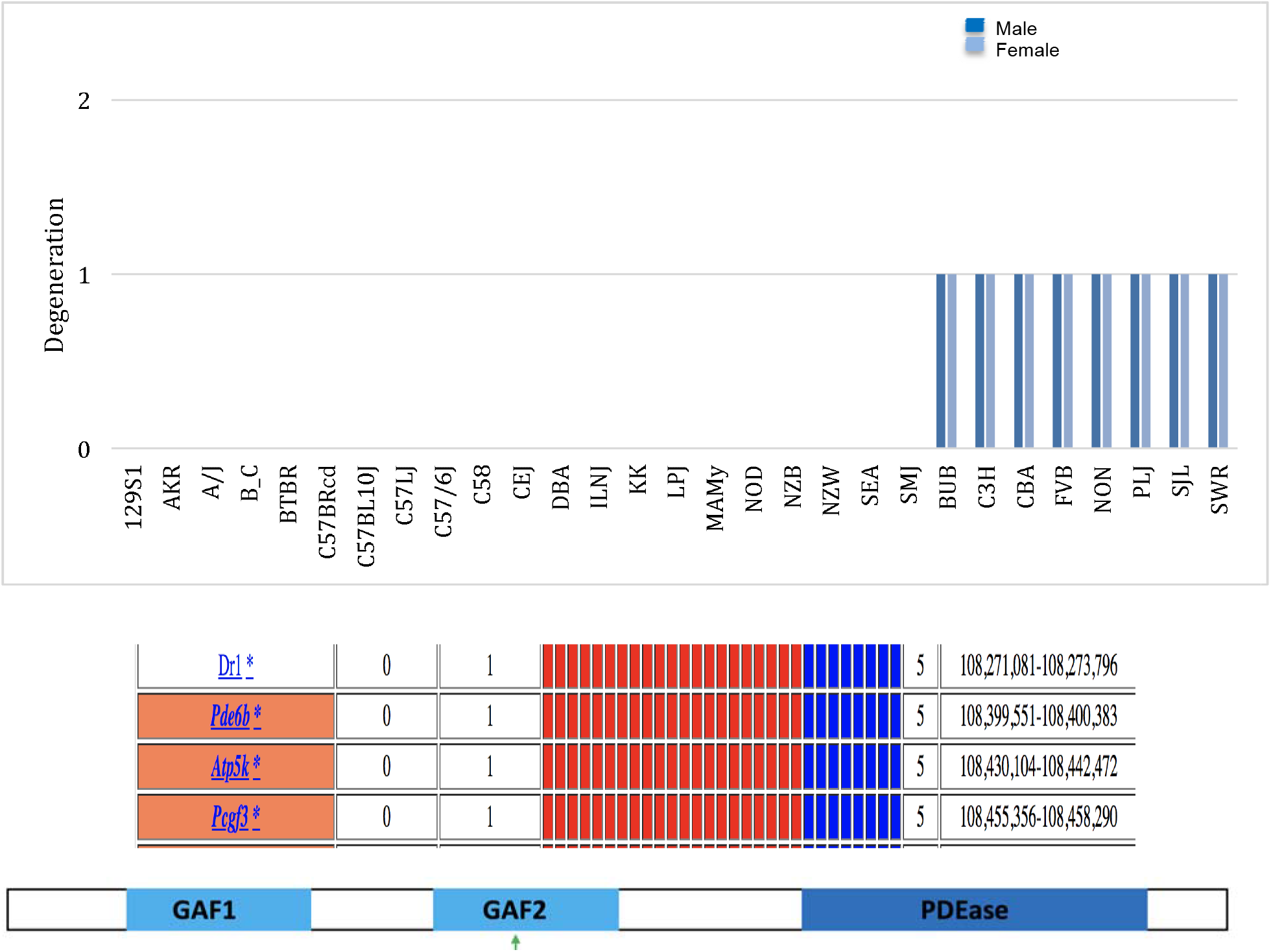
**Top:** The incidence of retinal degeneration in 29 inbred strains. Eight strains had significant retinal degeneration in all male and female mice (blue bars) examined, while 21 strains (indicated by the absence of a bar) had normal retinas. A bar with a value of 1 indicates all mice of that strain had retinal degeneration. **Middle:** HBCGM identified 33 genes with haplotype blocks whose allelic pattern was perfectly correlated with the retinal phenotype. However, only the four genes expressed in the retina are shown here. The genes within the correlated haplotype blocks are indicated by their symbol; and an orange, or white background indicates whether a SNP caused a significant or no amino acid change, respectively. The haplotypic pattern is shown as colored rectangles that are arranged in the same order as the input data shown above. Strains with the same colored rectangle have the same haplotype within the block. The p-values and genetic effect size were calculated as previously described [33]. **Bottom**: The domain structure of the Pde6b protein, and the location of its two GAF and the esterase domains are shown. The relative position of a SNP allele with a stop codon within the 2^nd^ GAF domain is indicated. The 8 strains with retinal degeneration all had the stop codon at position 347, while the 21 strains with normal retinas had the *Tyr347* allele.

Another data set (MPD: 9904) measured plasma high-density lipoprotein (**HDL**) cholesterol levels in female mice of 30 different inbred strains on a high-fat diet for 17 weeks [48]. There was a large inter-strain variation in HDL cholesterol levels (range 40 to 125 mg/dL) (**Fig. 3**), which was highly heritable (ANOVA p value = 3 × 10^−72^). Since this was a quantitative trait, many genes had haplotypic patterns that correlated with the HDL levels. Therefore, automated methods were used to evaluate the top 50 gene (p value = 4×10^−07^) candidates identified by the HBCGM analysis to identify genes that were: (i) expressed in liver, and (ii) contained a codon-changing SNP. Application of these criteria reduced the number of candidates to three genes (*Tomm40l, Nr1i3* and *Apoa2*), which were all located within a single genomic region (Chr 1, 171 MB), and a literature search indicated that only one was related to cholesterol metabolism. *Apoa2* encodes apolipoprotein (Apo-) A-II, which is the second most abundant protein within HDL particles, and it is involved in the lipoprotein metabolism pathway. *Apoa2* alleles were previously shown to affect HDL size and composition in mice [49]; and HDL levels in *Apoa2* knockout mice were decreased by 70% [50]. ApoA-II also plays an important role in cholesterol efflux; it modulates the interaction of HDL with lipid transfer proteins and enzymes [51]. Although the roles of the Pde6b in retinal degeneration, ApoA-II in HDL metabolism, and of Nrlp1b in the anthrax toxin response were previously known, these results demonstrate that genetic factors underlying important biomedical traits can be identified by HBCGM analysis of a large number of phenotypic datasets. However, candidate gene filtering methods were required for successful analysis of these traits, and these structured methods enabled the analyses to be much more rapidly completed. An example of a potential novel genetic finding for another retinal trait is described in the supplement (**Fig. S2**).

**Figure 3.**
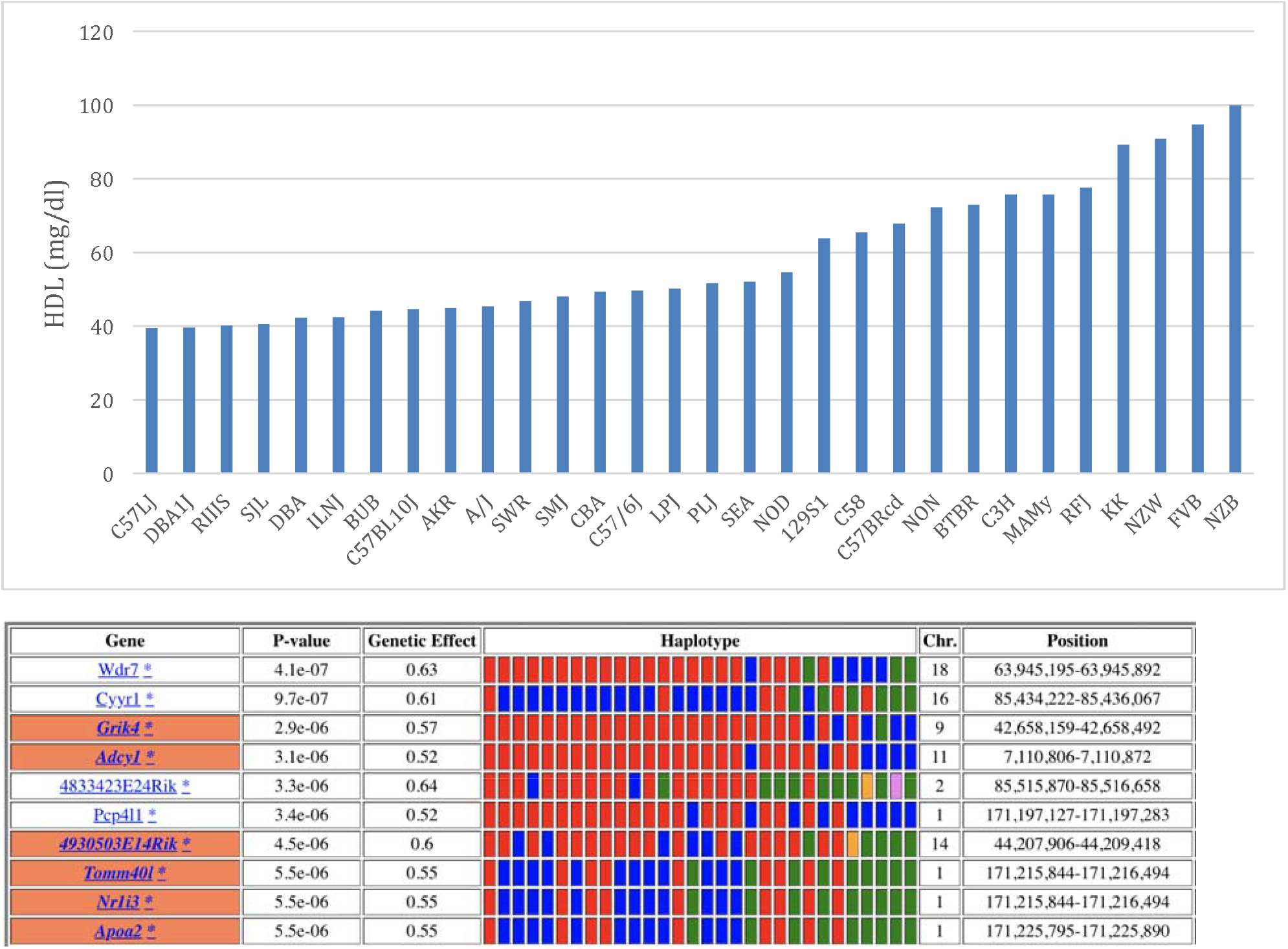
**Top:** Plasma HDL levels were measured in female mice of 30 inbred strains maintained on a high-fat diet for 17 weeks. Each bar is the average of the plasma HDL ± SEM (mg/ml) measured in female mice (n=5-14 mice per group) of the indicated strain. **Bottom:** The HBCGM program identified the ten genes whose allelic pattern was most highly correlated with plasma HDL levels. The genes within the correlated haplotype blocks are indicated by their symbol; and an orange or white background indicates whether SNPs cause a significant or no amino acid change, respectively. The haplotypic pattern is shown as colored rectangles that are arranged in the same order as the input data (shown in the graph above). Strains with the same colored rectangle have the same haplotype within the block. The p-values and genetic effect size were calculated as previously described [33]. Of note, since the calculated genetic effect size for the *Apoa2* alleles was 0.55, other genetic factors could also affect the plasma HDL levels in murine strains.

### A novel genetic effector mechanism

Another dataset (MPD: 50243) measured hepatic succinylcarnitine levels in 16-week-old male mice after a 16-hour fast [52]. The levels were highly variable and were highly heritable (ANOVA p value = 1×10^−18^) across the 24 strains. Two (FVB, SJL) of the 24 strains had a very high hepatic level of this metabolite (**Fig. 4**). Our own analysis confirmed that SJL mice had dramatically increased hepatic succinylcarnitine levels (**Fig. S3**). HBCGM analysis identified 152 genes whose allelic pattern correlated with the hepatic succinylcarnitine level (i.e. the FVB and SJL haplotype differed from the 22 other strains). However, only 3 of these genes were expressed in liver and had a codon changing SNP; and a literature search revealed that only one was linked with succinylcarnitine metabolism. *Lactb* encodes a serine beta-lactamase-like protein that forms filaments that localize to the region between the inner and outer mitochondrial membranes, and it has been proposed that Lactb filaments could affect mitochondrial organization and mitochondrial metabolism [53]. Moreover, analysis of two large population-based European cohorts revealed that human *LACTB* alleles were associated with plasma succinylcarnitine levels (rs2652822, p=7×10^−27^) [54], and the level of *Lactb* mRNA expression positively correlates with body mass index (*p*□= □1.19□× □10^−8^) [55]. Expression of a *Lactb* transgene also was shown to affect fat mass in mice [56, 57].

**Figure 4.**
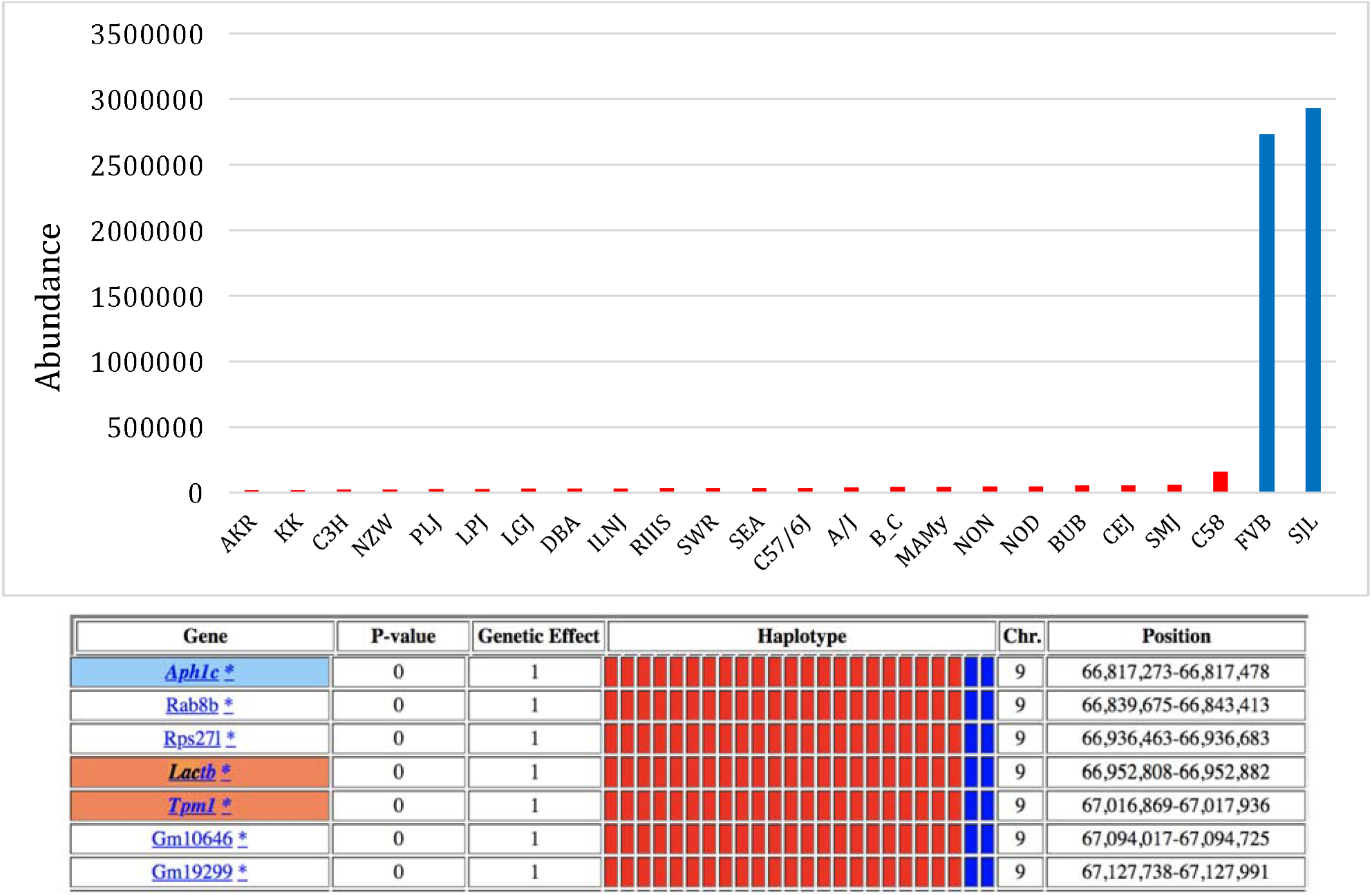
**Top:** Hepatic succinylcarnitine levels were measured in 16-week-old male mice after a 16-hour fast in 24 inbred strains. Each bar is the average succinylcarnitine level ± SEM (shown as the abundance determined by mass spectroscopy) for each strain. **Bottom:** HBCGM analysis identified 152 genes whose allelic patterns were highly correlated with the succinylcarnitine level. However, only 3 of these genes (colored gene symbol background) were expressed in the liver and had a SNP causing change in the predicted amino acid sequence. The genes within correlated haplotype blocks are indicated by their symbol: an orange, white, or blue background indicates whether a SNP does or does not cause an amino acid change, or if it affects a splice site, respectively, is present. The haplotypic pattern is shown as colored rectangles that are arranged in the same order as the input data. Strains with the same colored rectangle have the same haplotype within the block. The p-values and genetic effect size were calculated as previously described [33].

Examination of the allelic pattern shared by the two strains (FVB, SJL) with elevated hepatic succinylcarnitine levels suggested a potential genetic effector mechanism (**Fig. S4**). Nuclear DNA encodes all except 13 of the ∼1158 proteins required for the assembly and function of mitochondria [58-60]. These proteins are synthesized in the cytoplasm and are then transported into the mitochondria. Although there are various targeting mechanisms [61], most have NH2-terminal mitochondrial targeting sequences (**MTS**) that are enriched in hydrophobic and positively charged amino acids. Comparative analyses of yeast, mouse and human MTS have indicated that their length, sequence and net charge (between +3 and +6) are highly conserved [62]. The sequence conservation is due to the fact that the MTS must: form amphiphilic α-helices; interact with subunits of a mitochondrial surface protein for mitochondrial import; and must then be removed from the mature protein by a cleavage reaction that is performed by a very limited set of proteases [61, 63-67]. FVB and SJ/L mice share unique alleles at five SNP sites: *Arg110Gly* and *Pro88Leu* are within the NH2-terminal MTS; while three SNPs (*Val217Ala, Ala266Thr* and *Ser237Pro*) are within the sequence of the mature Lactb protein. Since two SNPs (*Pro88Leu, Arg110Gly*) introduce significant amino acid changes within the MTS (**Fig. 5**), the mitochondrial localization of Lactb could be altered in FVB and SJL mice. To investigate this, cDNAs encoding the C57BL/6 and FVB allelic forms of *Lactb* were expressed as EGFP fusion proteins in 293T cells (**Fig. 6A**). The C57BL/6 Lactb-EGFP fusion protein was highly expressed; it had a punctate expression pattern that overlapped with cellular mitochondria; and it differed from that of a control (EGFP only) protein that was expressed throughout the cytoplasm. In contrast, the FVB Lactb-EGFP fusion protein was expressed at a much lower level than the C57BL/6 protein (**Fig. 6B**). Importantly, the mRNAs for the two allelic forms of these fusion proteins were expressed at the same level after transfection (**Fig 6C**). Since these cDNAs were transcribed at similar rates, the protein expression differences must result from an allelic effect on a post-transcriptional process. To more precisely determine the basis for this allelic effect, FVB alleles at two positions (88Leu, 110Gly) within the MTS were engineered into the C57BL/6 *Lactb* cDNA by site directed mutagenesis (Fig. 6A). Interestingly, the C57BL/6^88L 110Gly^ Lactb-EGFP fusion protein was expressed at a reduced level, which was similar to that of the FVB allelic form (**Fig. 6C**). Similar differences in protein expression were also noted when the different allelic forms of the fusion proteins were expressed in HepG2 cells (**Fig. S5**). These results indicate that while the C57BL/6 protein was efficiently expressed and transported into mitochondria, FVB alleles at two sites within its MTS dramatically reduced its expression level within mitochondria.

**Figure 5.**
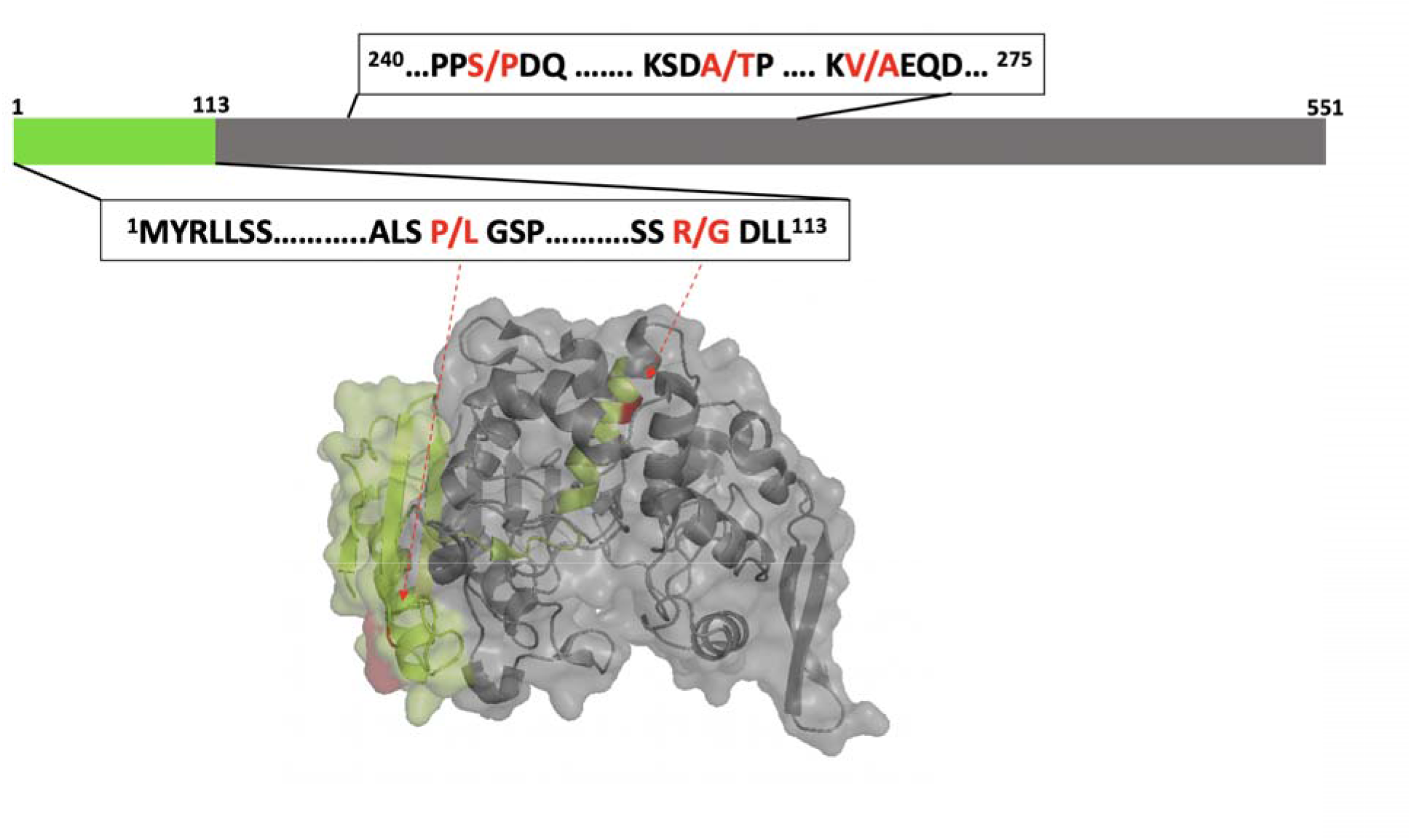
**Top:** Lactb protein domains and the sites of the SNP alleles. The green region is the 113-amino acid mitochondrial targeting sequence (MTS), and the grey region is the mature protein with the Lactb domain. The box below shows the position of two murine SNPs (*Pro88Leu* and *Arg110Gly*) within the MTS where the two strains with high succinylcarnitine levels (SJL, FVB) share unique alleles that are not present in other strains. The box above shows the position of three SNPs (*Ser247Pro, Ala266Thr, Val217Ala*) within the mature Lactb protein where the two strains (SJL, FVB) with high succinylcarnitine levels share unique alleles (*247Pro, Thr266, Ala271*). **Bottom**: A protein structural model of the Lactb protein was produced using I-TASSER [101]. The green region is the NH2-terminal MTS, and the gray region shows the structure of the mature protein. The positions of the two SNP sites within the MTS are shown in red.

**Figure 6.**
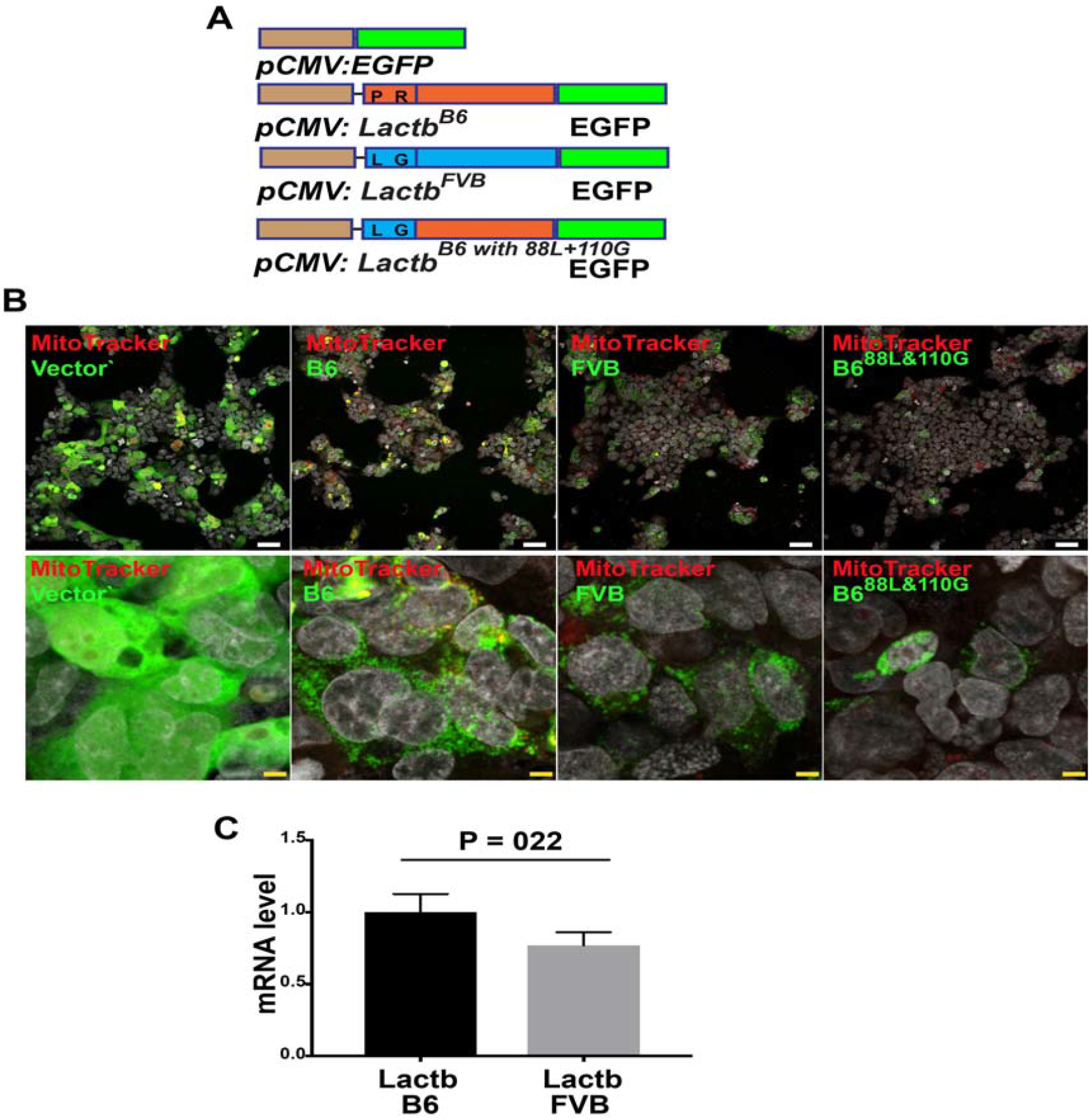
(**A**) Diagram of murine Lactb-EGFP fusion proteins. The control vector expresses an EGFP cDNA from a CMV promoter. The Lactb-EGFP fusion proteins were prepared from: (i) C57BL/6 or (ii) FBV *Lactb* mRNAs; or from a (iii) C57BL/6 *Lactb* mRNA with FVB alleles engineered at two positions (*88L, 110G*) within its MTS. (**B**) Confocal images obtained 24 hours after 293T cells were transfected with the plasmids shown in (A) indicate the different levels of expression and sub-cellular localization of the fusion proteins. For each construct, low magnification images are shown in the upper row (scale bar 50 um), and higher magnification images of a region within the upper panel (single cell level) are shown in the lower row (scale bar 5 um). Mitochondria are stained red, and the green color indicates the Lactb fusion protein expression. In contrast to the diffuse cytoplasmic expression pattern of the control EGFP protein, the C57BL/6 Lactb-EGFP fusion protein is expressed at a high level in a punctate pattern that overlapped with mitochondria. The FVB Lactb-EGFP protein was also expressed in a punctate pattern, but at a much lower level than the C57BL/6 Lactb-EGFP fusion protein. Also, the level and pattern of expression of the C57BL/6 ^88L 110G^ Lactb fusion protein resembled that of the FVB Lactb-EGFP fusion protein. This result is representative of 3 independently performed experiments. (**C**) RT-PCR measurement of the level of *Lactb-EGFP* mRNA expression in 293T cells 24 hrs after transfection with the plasmids shown in (A). Each bar is the average ± SE of 3 independent measurements. Despite the difference in the level of protein expression, there was no significant difference between the level of *C57BL/6* and *FVB Lactb-EGFP* mRNA expression (p=0.22).

### SNPs are present in the MTS of many murine and human proteins

To determine if allelic variation within the MTS could affect other nuclear-encoded mitochondrial proteins, we used our mouse SNP database to investigate whether MTS SNPs were present in any of the 524 genes that were annotated as nuclear encoded mitochondrial proteins [68]. We found 188 SNPs within the MTS of 120 of these murine genes; and 109 SNP alleles caused a major amino acid change in the predicted MTS of 79 genes (blosum-62 matrix score <-1) (**Table S2**). We then examined the NCBI SNP database to determine whether SNP alleles altered the MTS in 544 annotated human proteins [69], and found 161 codon-changing SNPs within the MTS of 83 of these proteins. It is noteworthy that 78 of these genes have other mutations that are located outside of the MTS, which cause human genetic diseases with very severe phenotypic effects (**Table S3**). Also, 8 genes have SNP alleles within their MTS that introduce a stop codon, and 12 genes have a SNP allele affecting the initiator methionine (**Table 1**). Moreover, allelic changes in 55 of these disease-associated proteins are predicted to have a major effect on the MTS sequence (blossom-62 matrix score ≤ -1). For example, there are 13 SNPs within the 33 amino acid MTS of a *thymidine kinase 2* (*TK2*) (**Table S3**). Genetic mutations that inactivate *TK2* cause mitochondrial DNA depletion, which presents in early childhood with a progressive myopathy or encephalopathy [70-73]. As another example, there were four cSNPs within the 53 amino acid MTS of *Pyruvate Dehydrogenase Complex Component X (PDHX)* (Table S3), which encodes the E3 ubiquitin ligase binding protein of the pyruvate dehydrogenase complex that catalyzes the rate-limiting step in aerobic glucose oxidation. A genetic deficiency of PDHX produces a life-threatening condition that causes developmental retardation [74]. Interestingly, the *Arg23Cys and Arg24Gly* SNP alleles have a major effect on the charge (the minor alleles reduce the charge from +6.1 to +4.0) and isoelectric point (from pH 12.6 to pH 10.74) of the PDHX MTS. Hence, there are multiple examples of allelic effects that are likely to impact the mitochondrial localization, and possibly the function, of human proteins that are of importance for cellular and tissue function.

**Table 1.**
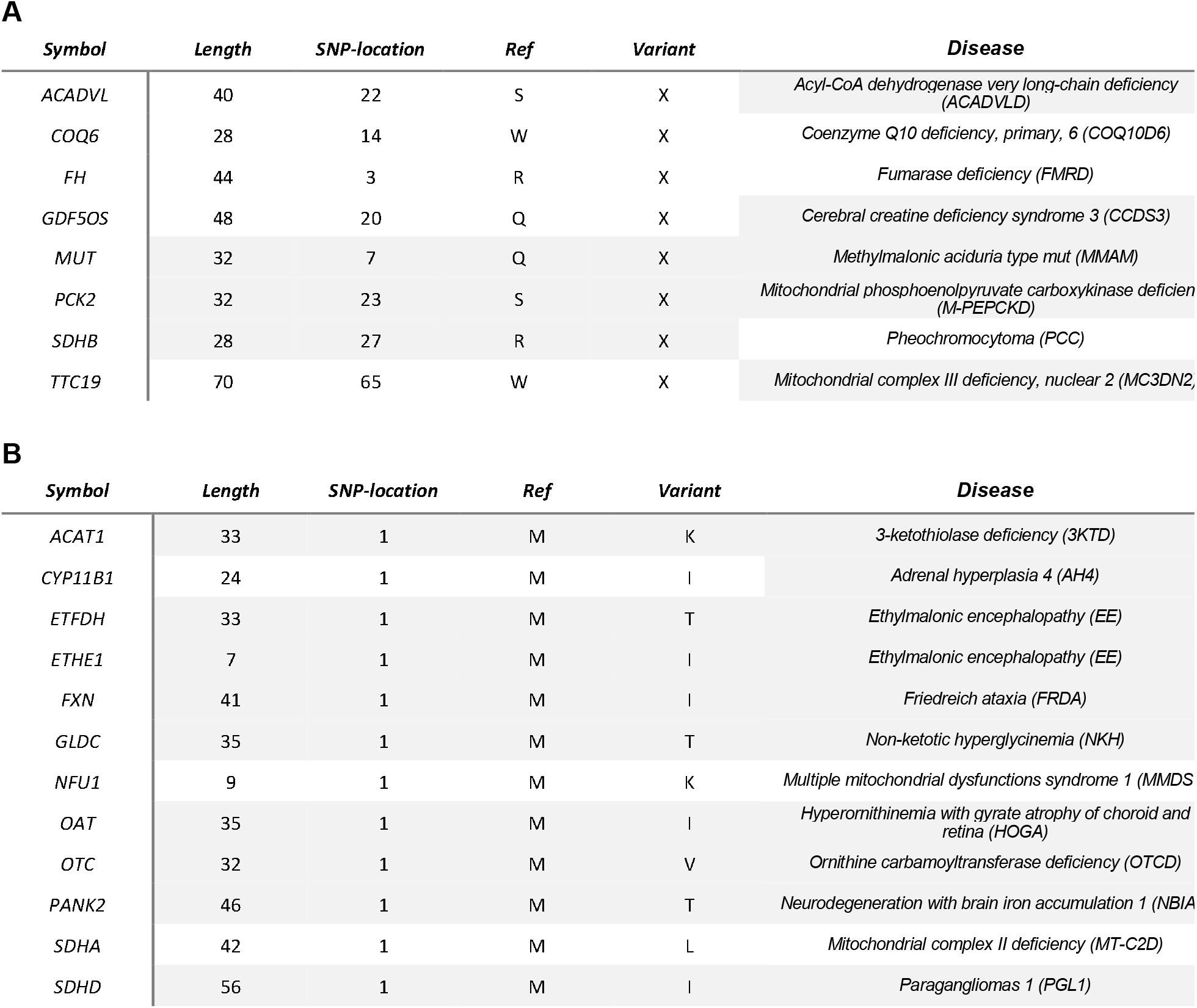
We used the NCBI human SNP database to identify SNP alleles that altered the MTS of the 544 genes that encoded human proteins that were annotated as having a MTS. We found human SNP alleles that introduced a premature termination codon (PTC) within the MTS of eight of these genes (**A**), and human SNPs that altered the initiator methionine for 12 genes (**B**). This table shows the gene symbol, the predicted length of the amino-terminal MTS, the position of the altered amino acid within the MTS, and the identity of the reference and variant amino acid for each SNP. These human SNPs are of interest because they are located within genes that have one or more other mutations, which are located outside of the MTS, that have been shown to cause a human genetic disease. The name of the disease caused by other SNPs (outside of the MTS) within each indicated gene (obtained from the Mendelian Inheritance in Man database) is shown. While the MTS SNPs are not disease associated, they could be of interest. For example, seven of the 8 genes, which have a SNP that introduces a PTC allele in their MTS are associated with metabolic changes that cause severe diseases. Thus, an individual that expresses a truncated form of any of the proteins encoded by these genes could have a metabolic abnormality. Similarly, a polymorphism affecting the initiator methionine of any of the 12 genes shown could have an impact on several phenotypes.

## Discussion

We demonstrate how computational analysis of a large biomedical response database could accelerate the pace of genetic discovery. Our ability to analyze this large phenotypic database is dependent upon two factors: (i) an increased number of inbred strains whose genome has been sequenced; and (ii) the use of automated methods for analyzing HBCGM output. *Why is the breadth of strain coverage important?* The database analyses responses across many strains, and this differs from prior mouse genetic analysis methods. For many years, mouse genetic models were analyzed by characterizing intercross progeny generated from two parental strains. This required a large amount of time for the generation and analysis of intercross progeny, and causative genetic factors could not be precisely localized [75]. To improve genetic mapping precision [76, 77], investigators have produced large panels of recombinant inbred strains (i.e. increased depth), but the progeny are generated from only 6-8 founder strains [78, 79]. While the increased depth can improve genetic mapping precision, its utility is limited by the lack of strain breadth. When a small number of strains are evaluated, the actual extent of phenotypic variation that is present in the mouse population is under-estimated [10, 34]. This is a critical, since a key factor for successful genetic discovery is analyzing strains that exhibit outlier responses. For example, the MTS allelic effect on mitochondrial metabolism could not have been uncovered using any of the available recombinant inbred strain panels [78, 79] since the two strains (SJL, FVB) with high hepatic succinylcarnitine levels were not among the founder strains used to generate the panel. In fact, our initial analysis of the 8462 MPD datasets indicated that inbred strains exhibiting outlier responses (i.e. those in the top or bottom 10%) were often not found among the 23 strains previously in our SNP database [15]. The number of evaluable datasets increased when the genome sequence of 43 strains became available. Each inbred strain has unique genetic variants, and possibly phenotypic responses, which could enable genetic discovery. Our projections [10] indicate that over ∼100,000 new SNPs per strain will be found even after the genomes of 40 strains are sequenced. As the emphasis in 21^st^ Century healthcare shifts from disease treatment to disease prevention [80], new murine genetic models will be needed for the new phenotypes that will be of interest 10 or more years from now. Since we cannot predict which strains will have outlier responses for phenotypes of future interest, obtaining the genomic sequence for an increasing number of the >450 available inbred strains [81] is of great importance for 21^st^ Century genetic discovery.

*Why are automated and structured methods needed for GWAS data analysis?* Filtering methods are required for selecting a true causative factor from among the many genomic regions that correlate with a phenotypic response pattern. On multiple prior occasions [36], we have found that causative genetic factors could be identified when other types of data were used to filter the gene candidates output by HBCGM analysis [13, 22, 24, 25, 37]. Since those analyses were manually performed, and they examined one dataset at a time, this filtering process is far too cumbersome for analyzing the 8462 available MPD datasets. Therefore, to select the most likely gene candidates among those output by HBCGM analysis, we developed selection criteria that could be automated. In these initial studies, we utilize gene expression criteria, select candidate genes with cSNPs; and use available information in published literature to identify the most likely causative gene(s) output by the genetic mapping program. This resembles methods developed by others that use: transcriptome wide association results [82, 83] or functional information [84-86] to select causative loci from among the many SNP sites identified in a human GWAS; or those that identify SNPs near *a priori* identified trait-related gene candidates in plant GWAS analyses [87]. The automated analysis of 152 correlated genes in the HBCGM output for the hepatic succinylcarnitine data led to the identification of four likely candidate genes, which was quickly narrowed to one obvious candidate by the literature search. While the criteria used in our initial studies select for cSNPs, a recent analysis of developmental disorders [88] indicated that allelic variation within non-coding regions could impact many other traits. We anticipate that filtering process improvements will enable other types of SNPs to be identified. Improved methods for analyzing the impact of allelic changes in non-coding sequences [89] could subsequently be used. A method for automated identification of genetic factors underlying metabolomic differences [90] could enable metabolomic data to be incorporated into the analyses. Our rudimentary literature searches could be improved by using a deep neural network [91], which has already been used to identify mutations that cause rare diseases in human populations. Implementation of these methods could enable ‘augmented intelligence’ further improve genetic discovery capabilities. This could enable the computational power and the large number of available phenotypic datasets to be used to advance our understanding of how biomedical traits are genetically regulated.

These computational methods identified a novel genetic effector mechanism: allelic changes within the MTS of murine *Lactb* alter its expression and mitochondrial localization. Moreover, this genetic effector mechanism could be active in other murine and human nuclear encoded mitochondrial proteins, which suggests that it could be of broad importance for disease susceptibility. In addition to the many known genetic diseases that are associated with nuclear encoded mitochondrial proteins [92], mitochondrial dysfunction is fundamental to many commonly occurring disease [93, 94] and age-associated conditions [59], with AIDs progression [95], and with cancer susceptibility [96]. In one case, *glutathione peroxidase 1* (*Pro198Leu)* alleles were shown to differentially affect its relative expression level in mitochondria, and altered the cellular response to oxidative stress [97]. Detailed studies in mice [98] and fish [99] have demonstrated that polymorphisms within the mitochondrial and nuclear genomes interact, and these interactions affect physiologically important processes. However, little was known about the effect of MTS polymorphisms on phenotypic responses and/or disease susceptibility. Of particular importance, MTS SNPs are present in many human proteins, including those where mutations outside of their MTS have caused very genetic diseases with a very severe impact. Since mutations within these genes have such significant health consequences, it is likely that at least some of the MTS allelic changes will impact other biomedical traits and possibly disease susceptibility.

## Conclusions

Implementation of the computational analysis methods described here could enable augmented intelligence to be used for genetic discovery. This could enable the computational power and the large number of phenotypic datasets that are now available to be used to advance our understanding of how biomedical traits are genetically regulated.

Methods are available in the online supplement.

## Data availability

The data sets within the Mouse Phenome Database (**MPD**) that were analyzed in this study are available at (https://phenome.jax.org). All sequence data is available at http://www.ncbi.nlm.nih.gov/bioproject/593371 (Bioproject ID: PRJNA593371). The HBCGM output for all of these datasets is available https://drive.google.com/file/d/1ryL_R0DKN4a_414BS1uCS2-S5bwCtC/view?usp=sharing. The source code for candidate gene filtering is available at http://github.com/AhmedArslan/HbCGM_paper, and that used for the functional characterization of genes is available at [100].

## Acknowledgements

We thank Dr. Robert Lewis for helpful comments and for reviewing this manuscript.

## Author contributions

GP and AA wrote the paper; AA, MZ, XC, WZ, MF, and RD generated data; and AA, MZ, DD, FZ and GP analyzed the data.

### Abbreviations

cSNP: codon-changing SNP
GWAS: genome-wide association study
HBCGM: haplotype-based computational genetic mapping
MTS: Mitochondrial targeting sequence.

