## Supplementary material for "High Throughput Computational Mouse Genetic Analysis": suppl info: Supp Info v7.docx

**Materials and Methods**

*Genomic sequencing and analysis*. The genomic sequence of 33 inbred strains was determined as previously described [1]. The genomic sequence of the following 20 additional strains was generated using DNA obtained from the Jackson Laboratory (Bar Harbor, ME): CE/J, DBA/1J, KK/HlJ, NON/ShiLtJ, NU/J, P/J, PL/J, RF/J, RHJ/LeJ, RIIIS/J, NZW/LacJ, NZB/BINJ, FVB/NJ, BUB/BnJ, NOR/LtJ, TALLYHO/JngJ, RBF/DnJ, BPL/1J, BPN/3J, MRL/MpJ. The sequence data was analyzed as previously described [1]. In brief, the sequencing reads were filtered and trimmed, then the resulting reads were aligned to the reference genome (C57BL6/J, mm10) using Burrow-Wheeler Aligner (BWA) [2] .

For comparison purposes, we also performed base quality score recalibration, indel realignment, and duplicate removal using GATK (v4.1.5). Joint variant calling across 53 inbred strains were performed using BCFtools (v1.9) [3] or GATK HaplotypeCaller [4] using their default settings. To ensure high quality SNV calls, we only selected single nucleotide variants using the following parameters: QUAL scores >= 50, the GT value must be a homozygous alternative call and the Phred-scaled heterozygous genotype likelihood (PL) value is at least 20 units above that of any other homozygous PL value. A whole genome SNP database with 23.6M SNPs and alleles covering 53 inbred strains was produced using BCFtools pipeline.

To assess the variant calls, SNP alleles were downloaded from dbSNP (build 142) via the mouse genome informatics (MGI) website (<http://www.informatics.jax.org/snp>). We excluded SNPs from 4 strains (BALB/cJ, LP/J, NZB/B1NJ, SPRET/EiJ) because there were discrepancies between the MGI and Mouse Genome Project data (https://www.sanger.ac.uk/data/mouse-genomes-project/). For the analysis, a correct call is defined as one where a variant is detected and is the same as that in dbSNP. An incorrect call is one where a variant is called, but it is different from that in dbSNP; while missed call is one where a variant that is present in dbSNP is not identified.

*Haplotype block construction and genetic mapping*. HBCGM was performed as originally described [5] using the modifications described in [6]. In brief, HBCGM utilizes bi-allelic SNPs that are polymorphic among the strains. Only bi-allelic SNPs with at least one definitive homozygous alternative call among the strains were included, while all other variants (including INDELs) were marked and excluded from HT-HBCGM analysis. The location and potential codon-change caused by a SNP are then annotated relative to the encoded gene using predictive gene models from Ensembl version 65. Only SNPs meeting the following criteria were used for haplotype block construction: (i) polymorphic among the strains with input trait data; and (ii) there were at least 8 strains (or one half of the number of input strains) with unambiguous allele calls. Haplotype blocks with 2, 3, 4 or 5 haplotypes were then dynamically produced as described in, and the correlation between the input phenotypic data and the haplotype pattern within each identified block was evaluated as described [6], The genes are then sorted based upon the ANOVA p value (in increasing order) for numeric data or by the F statistic (in decreasing order) for categorical data. For a gene that is covered by multiple blocks, the smallest p value (or the largest F statistic) obtained for all blocks within that gene was used.

The genetic effect size (η^2^) is calculated using our previously described method: [1] [7]

$$\eta\text{2}=\frac{\sigma\text{2}\text{B}}{\sigma\text{2}\text{T}}=\frac{SSB}{SST}$$

where SSB is the between-group sum-of-squares of the ANOVA model given as and SST is the total sum-of-squares. η^2^ is the genetic effect of the groups defined by haplotypes on the trait value and the total variance ($\sigma\text{2}\text{T}$) consists of within-group variance and between-group variance given as:

$$\sigma\text{2}\text{T = }\sigma\text{2}\text{B + }\sigma\text{2}\text{w}$$

for a sample size of n with *k* groups, with equal group sizes the *F* statistics of samples with effect size $\eta\text{2}$ follows a noncentral *F* distribution as *F*(k – 1, n – k, λ) with the noncentrality parameter:

$$\lambda=n\sigma\text{2}\text{B}/\sigma\text{2}\text{w}=n\sigma\text{2}\text{B}/n\sigma\text{2}\text{T}-n\sigma\text{2}\text{B}=n\eta\text{2}/1-\eta\text{2}$$

Therefore, the significance level α for power of one-way ANOVA test is given as:

Power(α,η^2^ ,n, k) = Prob(*F*(*k*-1,n-*k*,λ)) < *F* _crit_)

where *F* _crit_ = *F* _(1−α, k–1, n–_*_k_*_)_ is the (1−α) quantile of the *F* distribution with *k* – 1 and n – *k* degrees of freedom. Additional details about the HBCGM method are described elsewhere [5] [10] [7].

*HT-HBCGM analysis*. Given the large number of datasets within the MPD database (2138 after filtering for traits that have a strong genetic component), we elected to parallelize our algorithms. The haplotype blocks for a set of strains under analysis were constructed on a per-chromosome basis. Since there are 20 chromosomes (excluding the Y chromosome) in mouse, the computational time was reduced by a factor of 20. The permutation testing was performed in batches of up to 64 threads, reflecting the 64 processor cores we had available. The full analysis (association tests and calculation of false discovery rate p value) utilized 10 days of 64 processor cores.

*Filtering methods used to analyze genetic mapping output*. To identify likely causative genetic factors, a series of automated programs were used to analyze the genes with allelic patterns that were highly correlated with a phenotypic response pattern. The output genes were selected based upon the following criteria: (i) expression within an organ that is associated with the evaluated phenotype; (ii) functional evaluation of the effect of SNP alleles, which included selecting those that altered the predicted amino acid sequence of a protein; (iii) and a reported gene function based upon GO database annotation (i.e. enzyme, transcription factor, etc.) or a search of the published literature. A set of programs were established to perform the supervised filtering according, which provided an efficient method for identifying causative genetic factors in HBCGM output. To determine whether a gene was expressed within a target organ, preprocessed gene expression datasets were downloaded from the following sources: MGI [8], Expression Atlas [9] and Bgee [10]. Candidate genes were identified as expressed within a selected tissue using the predetermined expression threshold set by each resource. For example, using the parameters set by the Expression Atlas: a transcript per kilobase million (TRM) value >35 for a gene is considered as having a high expression level. Genes with TRM>35 within a target tissue were selected for further analyses. We next determined whether a SNP within a selected gene had a major effect the protein sequence. To do this, cSNP alleles that caused an amino acid to alternate between any of the following 4 groups were prioritized: non-polar and non-ionizable (A,V,L,I,P,W,F,M), polar and non-ionizable (G,S,T,Y,C,Q,N), basic (R,H,K), and acidic (D,E). To determine if cSNP alleles could affect protein structure or were within key functional regions of a protein, protein structure/function data obtained from several public resources was analyzed. The SNPs that overlapped with any protein functional or structural domain could have a major effect. SNPs within known protein structural domains (e.g. alpha helix, beta-sheets) were prioritized for further analyses. Protein secondary and tertiary structural information was assessed using the Protein Data Bank [11] information, and UniProt [12] and Ensembl databases were used to assess protein functional domains. To determine if a SNP was within protein functional or stability determining regions, including those affecting post translationally modified residues, protein annotation information in the following databases were analyzed: Phosphsite [13] and db PTM [14] databases were used to determine whether a SNP allele affected amino acid modifications (phosphorylation); and assessment of the SNP allele impact on protein functional domains was performed using data in the InterPro [15], Pfam [16] and SMART [17] databases. The Genome Vista [18] and Ensembl Genomes [19] databases were analyzed to determine whether a SNP was within a known promoter or enhancer region; and Metacyc [20] information was used to determine if a gene was within a metabolic pathway. For some datasets, the final analysis step was a natural language processing-based search [21] of the biomedical literature (PubMed) to determine whether a gene was linked with the input phenotype, or with a human or mouse disease. This search produced summary text, which highlighted the role of the gene relative to an input phenotype or disease process. A series of programs were developed to perform each of these analyses.

*LCMS measurement of succinylcarnitine abundance in liver*. The animal experiments were performed according to a protocol that was approved by the Stanford Institutional Animal Care and Use Committee. Liver tissue was obtained from 8-week old male SJL and B6 mice (n=5 per group). Food was withheld from the mice for 16 hours prior to sacrifice. The livers were weighed and re-suspended in 75% cold methanol and homogenized with a Precelly 24 Dual homogenizer (Bertin, France). Homogenate aliquots were centrifuged at 12K RPM for 10 minutes, and the supernatants were transferred to clean tubes. Succinylcarnitine derivatization with o-benzylhydroxylamine was carried out according to Tan et al [22]. In brief, o-benzylhydroxylamine and 1-ethyl-3-(3-dimethylaminopropyl)carbodiimide HCl (EDC) were added to the supernatants to achieve a final concentration of 0.2 M for both reagents. After room temperature incubation for 2 hours, the liquid was extracted into ethyl acetate, and the extracts were dried and re-suspended in 5% acetonitrile in 0.1% formic acid. LCMS analysis was carried out using an Agilent QTOF6520A coupled with UHPLC Infinity 1290 (Agilent, Santa Clara US). For chromatographic separation, a Phenomenex (Torrance, California, USA) Kinetex reverse-phase C18 column (dimension 2.1mmx100mm, 2.6mm particles, 100A ° pore size) was used; and solvent A was 0.1% formic acid in HPLC water and B was acetonitrile. Succinylcarnitine in the experimental samples was identified by comparing the retention time and accurate mass to that of a chemical standard (Sigma, St. Louis MO).

*Lactb structural model*. The structural model for the Lactb protein was produced using I-TASSER [23], which uses the information obtained from a database of experimentally resolved protein structures to generate a predicted structure. The default program settings were used for this model, and the input Lactb amino acid sequence was retrieved from UniProt database (ID: Q9EP89) [12].

*Lactb in vitro transfection studies*. *Lactb* cDNAs were generated by RT-PCR from liver tissue obtained from C57BL/6 and FVB mice. To produce the EGFP fusion protein, the full-length *Lactb* cDNAs were cloned (in frame) into the BamHI of pEGFP-N1 using the *NEBuilder* HiFi DNA Assembly (Ipswich, MA) system. Site-directed mutagenesis was used to introduce the *88L* and *110G* alleles into the C57Bl/6 *Lactb* cDNA, and this was performed using the NEBuilder HiFi DNA Assembly system. The sequences of all cDNAs were confirmed by Sanger sequencing. HepG2 cells (obtained from ATCC) were cultured in DMEM with 10% FBS. Then, 4 ug of each plasmid was transfected into wells with 1x10^6^ HepG2 cells using Lipofectamine (Invitrogen), and the cells were incubated for 24 hrs before analysis. The cells were then incubated with Mitrotracker (Invitrogen) at a 1:2000 dilution for 15-30 min, followed by fixation with paraformaldehyde and stained with DAPI. Images were obtained using Leica DMi8 or Leica TCS SP8 microscopes.

**Table S1:** The table shows the names of 43 inbred strains, the fold-coverage of their genomes (FC), and the number of SNPs identified (relative to the reference C57BL/6J strain).

| STRAIN # | STRAIN NAME | FC | SNP # |
| --- | --- | --- | --- |
| 1. | 129P2/OlaHsd | 35 | 5291668 |
| 2. | 129S1/SvImJ | 26 | 5214272 |
| 3. | 129S5SvEvBrd | 19 | 4629772 |
| 4. | A/J | 24 | 5062452 |
| 5. | AKR/J | 35 | 5098147 |
| 6. | BALB/cJ | 22 | 4617417 |
| 7. | B10 | 21 | 363118 |
| 8. | BTBRT<+> Itpr3<tf>/J | 19 | 4534249 |
| 9. | BUB/BnJ | 18 | 5146084 |
| 10. | C3H/HeJ | 29 | 5137933 |
| 11. | C57BL/10J | 24 | 336600 |
| 12. | C57BR/cdJ | 37 | 2598541 |
| 13. | C57L/J | 33 | 2479910 |
| 14. | C58/J | 42 | 2842374 |
| 15. | CBA/J | 24 | 5277532 |
| 16. | CE/J | 35 | 5651372 |
| 17. | DBA/2J | 22 | 5207181 |
| 18. | DBA/1J | 37 | 5253007 |
| 19. | FVB/NJ | 24 | 4914368 |
| 20. | I/LnJ | 32 | 5544345 |
| 21. | KK/HlJ | 37 | 6008628 |
| 22. | LG/J | 21 | 4988047 |
| 23. | LP/J | 19 | 5474423 |
| 24. | MA/MyJ | 21 | 4546167 |
| 25. | MRL/Mp | 27 | 4926824 |
| 26. | NOD/ShiLtJ | 23 | 5080934 |
| 27. | NON/ShiLtJ | 38 | 4995230 |
| 28. | NU/J | 39 | 4816135 |
| 29. | NZB/BlNJ | 27 | 5569902 |
| 30. | NZO/HlLtJ | 22 | 5349055 |
| 31. | NZW/LacJ | 26 | 5832209 |
| 32. | P/J | 40 | 5293988 |
| 33. | PL/J | 31 | 5286323 |
| 34. | PWK/PhJ | 23 | 10232623 |
| 35. | RF/J | 31 | 5312431 |
| 36. | RHJ/LeJ | 36 | 3927449 |
| 37. | RIIIS/J | 37 | 5284808 |
| 38. | SEA/GnJ | 40 | 4612347 |
| 39. | SJL/J | 43 | 5021350 |
| 40. | SM/J | 22 | 5219410 |
| 41. | ST/bJ | 64 | 5164286 |
| 42. | SWR/J | 20 | 4870923 |
| 43. | C57BL/6J | Ref | Ref |
| 44. | BPL | 62 | 4824215 |
| 45. | BPN | 76 | 4544195 |
| 46. | MOLF | 32 | 9467667 |
| 47. | CAST | 42 | 8269422 |
| 48. | NOR | 100 | 4860171 |
| 49. | PWD/PhJ | 137 | 10153715 |
| 50. | RBF | 168 | 6606273 |
| 51. | SPRET | 52 | 7848628 |
| 52. | TALLYHO | 134 | 5494776 |
| 53. | WSB | 39 | 5764665 |

**Table S2.** Murine genes with SNPs altering amino acids within mitochondrial targeting sequences (MTS). There are 188 SNPs that alter an amino acid with the MTS of 120 murine genes. This table shows the gene symbol, the length of the amino-terminal MTS, the location of the altered amino acid, and the identity of the reference and variant amino acid for each SNP.

| ***Symbol*** | ***Length*** | ***SNP-location*** | ***Ref*** | ***Variant*** |
| --- | --- | --- | --- | --- |
| *Aars2* | 23 | 15 | A | S |
| *Abcb10* | 82 | 79 | C | S |
| *Abcb10* | 82 | 52 | S | P |
| *Abcb8* | 38 | 19 | S | L |
| *Abhd10* | 43 | 25 | A | P |
| *Acaa2* | 16 | 10 | V | I |
| *Acadl* | 30 | 22 | L | P |
| *Acadl* | 30 | 14 | S | R |
| *Acsm2* | 46 | 25 | M | R |
| *Acss3* | 29 | 9 | R | C |
| *Afg3l1* | 70 | 61 | K | R |
| *Agmat* | 36 | 24 | A | E |
| *Agps* | 45 | 1 | M | L |
| *Agxt* | 23 | 5 | L | F |
| *Aldh4a1* | 23 | 18 | G | S |
| *Amt* | 28 | 4 | I | T |
| *Atpaf1* | 54 | 45 | L | I |
| *Auh* | 42 | 16 | G | R |
| *Auh* | 42 | 15 | V | G |
| *Auh* | 42 | 14 | T | A |
| *Bckdhb* | 48 | 43 | T | A |
| *Chdh* | 34 | 4 | V | A |
| *Chdh* | 34 | 12 | W | C |
| *Chdh* | 34 | 24 | Q | R |
| *Clpp* | 52 | 27 | H | R |
| *Clpx* | 56 | 37 | V | M |
| *Clybl* | 20 | 9 | T | A |
| *Cmpk2* | 73 | 57 | H | Y |
| *Coq2* | 34 | 15 | V | M |
| *Coq3* | 86 | 20 | F | L |
| *Coq4* | 30 | 14 | R | L |
| *Coq8a* | 159 | 137 | R | W |
| *Coq8a* | 159 | 159 | S | F |
| *Coq8a* | 159 | 103 | S | A |
| *Coq8a* | 159 | 56 | G | R |
| *Cox18* | 63 | 22 | G | R |
| *Cox8b* | 24 | 24 | H | Y |
| *Cpox* | 98 | 62 | G | R |
| *Cpox* | 98 | 66 | R | G |
| *Cyp11a1* | 36 | 30 | S | T |
| *Cyp24a1* | 35 | 34 | V | A |
| *Dars2* | 46 | 9 | R | S |
| *Dbt* | 61 | 29 | A | V |
| *Dglucy* | 26 | 13 | S | G |
| *Diablo* | 53 | 15 | Q | H |
| *Dlat* | 85 | 69 | R | S |
| *Dnlz* | 53 | 24 | R | Q |
| *Echdc2* | 17 | 14 | S | P |
| *Ecsit* | 48 | 5 | S | L |
| *Exog* | 41 | 39 | F | L |
| *Ftmt* | 49 | 7 | F | L |
| *Ftmt* | 49 | 13 | S | G |
| *Fxn* | 40 | 5 | G | R |
| *Gcdh* | 44 | 18 | R | Q |
| *Gcdh* | 44 | 9 | R | Q |
| *Gcsh* | 45 | 21 | T | A |
| *Gpd2* | 42 | 26 | P | Q |
| *Grpel2* | 31 | 21 | M | V |
| *Gsr* | 26 | 26 | A | T |
| *Hint2* | 17 | 13 | A | V |
| *Hspa9* | 46 | 21 | P | S |
| *Iba57* | 94 | 87 | T | A |
| *Iba57* | 94 | 86 | T | A |
| *Iba57* | 94 | 43 | P | S |
| *Iba57* | 94 | 36 | G | R |
| *Ifi27l2b* | 90 | 84 | T | A |
| *Ifi27l2b* | 90 | 80 | V | A |
| *Ifi27l2b* | 90 | 74 | I | T |
| *Ifi27l2b* | 90 | 69 | L | P |
| *Ifi27l2b* | 90 | 68 | V | A |
| *Ifi27l2b* | 90 | 59 | L | M |
| *Ifi27l2b* | 90 | 58 | G | S |
| *Ifi27l2b* | 90 | 54 | I | V |
| *Ifi27l2b* | 90 | 6 | V | L |
| *Ifi27l2b* | 90 | 5 | F | L |
| *Ifi27l2b* | 90 | 3 | R | K |
| *Lactb* | 113 | 110 | R | G |
| *Lactb* | 113 | 88 | P | L |
| *Lactb* | 113 | 23 | G | R |
| *Lrpprc* | 59 | 52 | R | L |
| *Mccc1* | 38 | 34 | A | V |
| *Mccc1* | 38 | 32 | P | L |
| *Mccc1* | 38 | 30 | H | R |
| *Mccc1* | 38 | 28 | T | M |
| *Mcee* | 38 | 18 | Q | X |
| *Mcee* | 38 | 29 | E | K |
| *Mcub* | 44 | 33 | D | G |
| *Mcub* | 44 | 21 | Q | R |
| *Mcub* | 44 | 5 | L | R |
| *Mdh2* | 24 | 23 | A | V |
| *Me2* | 18 | 10 | T | S |
| *Mettl17* | 19 | 17 | C | F |
| *Mgme1* | 64 | 30 | F | S |
| *Micu1* | 33 | 6 | T | A |
| *Micu2* | 22 | 10 | W | R |
| *Mmaa* | 62 | 31 | H | P |
| *Mmaa* | 62 | 12 | R | W |
| *Mrm2* | 24 | 23 | H | L |
| *Mrm2* | 24 | 10 | V | G |
| *Mrpl1* | 50 | 9 | R | C |
| *Mrpl1* | 50 | 19 | C | S |
| *Mrpl2* | 60 | 43 | P | H |
| *Mrpl2* | 60 | 51 | V | M |
| *Mrpl22* | 40 | 4 | A | T |
| *Mrpl3* | 40 | 19 | A | T |
| *Mrpl3* | 40 | 32 | I | T |
| *Mrpl30* | 34 | 8 | A | V |
| *Mrpl30* | 34 | 10 | P | Q |
| *Mrpl37* | 29 | 27 | K | R |
| *Mrpl42* | 31 | 11 | N | K |
| *Mrpl48* | 27 | 14 | W | R |
| *Mrpl55* | 32 | 19 | L | F |
| *Mrpl55* | 32 | 26 | L | F |
| *Mrps26* | 27 | 17 | P | R |
| *Mrps28* | 70 | 41 | D | A |
| *Mrps28* | 70 | 36 | E | K |
| *Mrps28* | 70 | 20 | L | F |
| *Mrps7* | 37 | 21 | C | Y |
| *Mrrf* | 55 | 16 | R | H |
| *Msrb2* | 42 | 7 | A | T |
| *Mterf1a* | 37 | 19 | D | G |
| *Mterf3* | 67 | 38 | R | H |
| *Mterf3* | 67 | 15 | K | N |
| *Mterf3* | 67 | 52 | S | C |
| *Mterf3* | 67 | 51 | T | A |
| *Mterf3* | 67 | 45 | P | A |
| *Mterf3* | 67 | 37 | V | E |
| *Mterf3* | 67 | 29 | R | H |
| *Mtif2* | 29 | 3 | Q | R |
| *Myg1* | 46 | 36 | P | L |
| *Naxe* | 53 | 38 | T | I |
| *Ndufb2* | 33 | 29 | S | G |
| *Ndufb5* | 46 | 16 | T | A |
| *Ndufb5* | 46 | 34 | T | M |
| *Ndufb8* | 28 | 21 | V | L |
| *Ndufs1* | 23 | 19 | V | A |
| *Ndufs2* | 33 | 8 | R | G |
| *Ndufs4* | 42 | 36 | Q | L |
| *Ndufs4* | 42 | 34 | L | F |
| *Ndufs4* | 42 | 25 | V | I |
| *Ndufs6* | 20 | 4 | V | A |
| *Ndufs6* | 20 | 3 | A | V |
| *Ndufs7* | 35 | 25 | R | H |
| *Nfu1* | 29 | 15 | V | A |
| *Nit1* | 33 | 22 | T | I |
| *Nlrx1* | 86 | 52 | Y | H |
| *Nlrx1* | 86 | 37 | F | V |
| *Nnt* | 43 | 35 | T | M |
| *Nubpl* | 38 | 6 | R | H |
| *Nubpl* | 38 | 33 | C | F |
| *Oxsm* | 27 | 19 | P | L |
| *Pars2* | 29 | 17 | C | Y |
| *Pck2* | 32 | 31 | H | R |
| *Phyh* | 30 | 2 | N | K |
| *Pink1* | 77 | 64 | S | L |
| *Pink1* | 77 | 63 | E | K |
| *Pink1* | 77 | 61 | M | I |
| *Pink1* | 77 | 13 | S | G |
| *Pink1* | 77 | 18 | Q | R |
| *Pink1* | 77 | 48 | Q | R |
| *Pisd* | 49 | 27 | R | L |
| *Pisd* | 49 | 23 | N | K |
| *Ptcd3* | 10 | 7 | A | V |
| *Rbfa* | 41 | 31 | S | A |
| *Rbfa* | 41 | 28 | L | P |
| *Rbfa* | 41 | 19 | W | R |
| *Rbfa* | 41 | 6 | A | V |
| *Rpusd4* | 46 | 29 | W | R |
| *Rsad1* | 17 | 12 | V | M |
| *Sdhc* | 29 | 10 | S | G |
| *Sdhc* | 29 | 4 | F | L |
| *Slc25a3* | 45 | 32 | P | S |
| *Slc8b1* | 26 | 20 | I | L |
| *Smdt1* | 47 | 14 | V | A |
| *Spg7* | 43 | 13 | P | H |
| *Spg7* | 43 | 18 | R | W |
| *Spg7* | 43 | 34 | S | L |
| *Stard7* | 61 | 45 | Y | H |
| *Suclg1* | 34 | 13 | T | I |
| *Tfb2m* | 43 | 12 | I | M |
| *Tmem65* | 55 | 26 | L | P |
| *Tmem70* | 77 | 72 | E | G |
| *Tmem70* | 77 | 74 | Q | R |
| *Top1mt* | 43 | 42 | S | N |
| *Txn2* | 59 | 44 | V | G |
| *Uqcrc1* | 34 | 11 | T | M |
| *Uqcrc1* | 34 | 24 | Q | R |
| *Yrdc* | 56 | 56 | P | A |

**Table S3.** Human genes with non-synonymous SNPs within mitochondrial targeting sequences (MTS). There are 161 SNPs that alter an amino acid with the MTS of 83 human genes. This table shows the gene symbol, the length of the amino-terminal MTS, the position of the altered amino acid within the MTS, and the identity of the reference and variant amino acid for each SNP. For the genes with mutations that are causative of human genetic diseases, the name of the genetic disease and the Mendelian Inheritance in Man (MIM) database identification number is shown.

| ***Symbol*** | ***Length*** | ***SNP-location*** | ***Ref*** | ***Variant*** | ***Disease*** | ***MIM*** |
| --- | --- | --- | --- | --- | --- | --- |
| *ACADM* | 25 | 6 | G | R | *Acyl-CoA dehydrogenase medium-chain deficiency (ACADMD)* | *201450* |
| *ACADM* | 25 | 4 | T | A |  |  |
| *ACADM* | 25 | 7 | E | K |  |  |
| *ACADM* | 25 | 17 | R | H |  |  |
| *ACADM* | 25 | 14 | L | F |  |  |
| *ACADM* | 25 | 9 | Q | R |  |  |
| *ACADSB* | 33 | 13 | R | K | *Short/branched-chain acyl-CoA dehydrogenase deficiency (SBCADD)* | *610006* |
| *ACADVL* | 40 | 17 | L | F | *Acyl-CoA dehydrogenase very long-chain deficiency (ACADVLD)* | 201475 |
| *ACADVL* | 40 | 22 | S | X |  |  |
| *ACAT1* | 33 | 1 | M | K | *3-ketothiolase deficiency (3KTD)* | *203750* |
| *ACAT1* | 33 | 3 | N | S |  |  |
| *ACAT1* | 33 | 32 | N | H |  |  |
| *ACAT1* | 33 | 26 | G | A |  |  |
| *ACSF3* | 83 | 17 | A | P | *Combined malonic and methylmalonic aciduria (CMAMMA)* | *614265* |
| *ACSF3* | 83 | 10 | R | W |  |  |
| *AIFM1* | 54 | 27 | N | S | *Combined oxidative phosphorylation deficiency 6 (COXPD6)* | 300816 |
| *AIFM1* | 54 | 21 | G | R |  |  |
| *ALAS2* | 49 | 4 | G | S | *Anemia, sideroblastic, 1 (SIDBA1)* | *300751* |
| *ALAS2* | 49 | 12 | K | Q |  |  |
| *APOPT1* | 39 | 27 | P | T |  |  |
| *AUH* | 67 | 16 | R | W | *3-methylglutaconic aciduria 1 (MGA1)* | *250950* |
| *BCKDHA* | 45 | 39 | P | H | Maple syrup urine disease 1A (MSUD1A) | 248600 |
| *BCKDHB* | 50 | 29 | A | V | Maple syrup urine disease 1B (MSUD1B) | 248600 |
| *BCKDHB* | 50 | 49 | V | G |  |  |
| *BCKDHB* | 50 | 31 | G | D |  |  |
| *CHCHD10* | 16 | 15 | R | L | *Frontotemporal dementia and/or amyotrophic lateral sclerosis 2 (FTDALS2)* | *615911* |
| *COQ2* | 34 | 28 | S | N | *Coenzyme Q10 deficiency, primary, 1 (COQ10D1)* | *607426* |
| *COQ4* | 30 | 20 | R | Q | *Coenzyme Q10 deficiency, primary, 7 (COQ10D7)* | 616276 |
| *COQ6* | 28 | 14 | W | X | *Coenzyme Q10 deficiency, primary, 6 (COQ10D6)* | 614650 |
| *COQ8A* | 162 | 85 | H | Q | *Coenzyme Q10 deficiency, primary, 4 (COQ10D4)* | *612016* |
| *CPOX* | 110 | 29 | Q | E | *Hereditary coproporphyria (HCP)* | *121300* |
| *CYC1* | 84 | 37 | W | C | *Mitochondrial complex III deficiency, nuclear 6 (MC3DN6)* | 615453 |
| *CYP11A1* | 39 | 31 | A | V | *Adrenal insufficiency, congenital, with 46,XY sex reversal (AICSR)* | *613743* |
| *CYP11B1* | 24 | 7 | P | Q | *Adrenal hyperplasia 4 (AH4)* | *202010* |
| *CYP11B1* | 24 | 1 | M | I |  |  |
| *D2HGDH* | 13 | 13 | I | S | *D-2-hydroxyglutaric aciduria 1 (D2HGA1)* | *600721* |
| *D2HGDH* | 13 | 2 | P | L |  |  |
| *DARS2* | 47 | 45 | S | G | *Leukoencephalopathy with brainstem and spinal cord involvement and lactate elevation (LBSL)* | *611105* |
| *DGUOK* | 39 | 21 | R | H | *Mitochondrial DNA depletion syndrome 3 (MTDPS3)* | *251880* |
| *DHODH* | 10 | 7 | K | Q | *Postaxial acrofacial dysostosis (POADS)* | 263750 |
| *DHODH* | 10 | 9 | G | R |  |  |
| *DIABLO* | 55 | 29 | S | L | *Deafness, autosomal dominant, 64 (DFNA64)* | *614152* |
| *DIABLO* | 55 | 53 | S | L |  |  |
| *EARS2* | 41 | 17 | G | C | *Combined oxidative phosphorylation deficiency 12 (COXPD12)* | 614924 |
| *EARS2* | 41 | 15 | R | W |  |  |
| *EARS2* | 41 | 3 | E | Q |  |  |
| *EARS2* | 41 | 41 | K | E |  |  |
| *ECHS1* | 27 | 2 | A | G | *Mitochondrial short-chain enoyl-CoA hydratase 1 deficiency (ECHS1D)* | 616277 |
| *ETFDH* | 33 | 31 | T | I | *Glutaric aciduria 2C (GA2C)* | *231680* |
| *ETFDH* | 33 | 1 | M | T | *Ethylmalonic encephalopathy (EE)* | *602473* |
| *ETFDH* | 33 | 23 | A | T |  |  |
| *ETHE1* | 7 | 1 | M | I |  |  |
| *FECH* | 54 | 24 | R | Q | *Protoporphyria, erythropoietic, 1 (EPP1)* | 177000 |
| *FH* | 44 | 25 | R | Q | *Fumarase deficiency (FMRD)* | *606812* |
| *FH* | 44 | 31 | N | T |  |  |
| *FH* | 44 | 3 | R | X |  |  |
| *FXN* | 41 | 1 | M | I | *Friedreich ataxia (FRDA)* | *229300* |
| *GATM* | 37 | 20 | W | L | *Cerebral creatine deficiency syndrome 3 (CCDS3)* | *612718* |
| *GDF5OS* | 48 | 20 | Q | X |  |  |
| *GDF5OS* | 48 | 42 | R | S |  |  |
| *GDF5OS* | 48 | 21 | R | Q |  |  |
| *GDF5OS* | 48 | 40 | R | C |  |  |
| *GLDC* | 35 | 1 | M | T | *Non-ketotic hyperglycinemia (NKH)* | *605899* |
| *GTPBP3* | 81 | 52 | R | Q | *Combined oxidative phosphorylation deficiency 23 (COXPD23)* | 616198 |
| *IARS2* | 48 | 14 | A | V | *Cataracts, growth hormone deficiency, sensory neuropathy, sensorineural hearing loss, and skeletal dysplasia (CAGSSS)* | 616007 |
| *IBA57* | 39 | 18 | G | S | *Multiple mitochondrial dysfunctions syndrome 3 (MMDS3)* | 615330 |
| *IDH2* | 39 | 10 | R | W | *D-2-hydroxyglutaric aciduria 2 (D2HGA2)* | 613657 |
| *ISCU* | 34 | 25 | G | E | *Myopathy with exercise intolerance Swedish type (MEIS)* | *255125* |
| *IVD* | 32 | 4 | M | L | *Isovaleric acidemia (IVA)* | 243500 |
| *L2HGDH* | 51 | 18 | L | R | L-2-hydroxyglutaric aciduria (L2HGA) | 236792 |
| *L2HGDH* | 51 | 10 | G | D |  |  |
| *MCCC1* | 41 | 31 | K | E | *3-methylcrotonoyl-CoA carboxylase 1 deficiency (MCC1D)* | *210200* |
| *MCCC1* | 41 | 8 | G | E |  |  |
| *MLYCD* | 39 | 3 | G | D | *Malonyl-CoA decarboxylase deficiency (MLYCD deficiency)* | *248360* |
| *MMAA* | 65 | 42 | Y | C | *Methylmalonic aciduria type cblA (MMAA)* | 251100 |
| *MRPL44* | 30 | 9 | L | P | *Combined oxidative phosphorylation deficiency 16 (COXPD16)* | *615395* |
| *MRPS16* | 34 | 12 | Y | H | *Combined oxidative phosphorylation deficiency 2 (COXPD2)* | 610498 |
| *MRPS7* | 37 | 2 | A | V | *Combined oxidative phosphorylation deficiency 34 (COXPD34)* | *617872* |
| *MUT* | 32 | 7 | Q | X | *Methylmalonic aciduria type mut (MMAM)* | *251000* |
| *NDUFA10* | 35 | 2 | A | G | *Leigh syndrome (LS)* | 256000 |
| *NDUFAF6* | 44 | 6 | R | G | *Mitochondrial complex I deficiency (MT-C1D)* | *252010* |
| *NDUFAF6* | 44 | 26 | A | P |  |  |
| *NDUFAF6* | 44 | 32 | I | K |  |  |
| *NDUFAF6* | 44 | 13 | I | T |  |  |
| *NDUFAF7* | 46 | 39 | P | T | *Defects in NDUFAF7 may be a cause of susceptibility to pathologic myopia* |  |
| *NDUFAF7* | 46 | 30 | H | R |  |  |
| *NDUFS1* | 23 | 5 | R | Q | *Mitochondrial complex I deficiency (MT-C1D)* | *252010* |
| *NDUFS2* | 33 | 20 | P | T | Mitochondrial complex I deficiency (MT-C1D) | 252010 |
| *NDUFV2* | 32 | 29 | V | G |  |  |
| *NFU1* | 9 | 1 | M | K | *Multiple mitochondrial dysfunctions syndrome 1 (MMDS1)* | *605711* |
| *NNT* | 43 | 33 | A | V | *Glucocorticoid deficiency 4 with or without mineralocorticoid deficiency (GCCD4)* | *614736* |
| *NUBPL* | 38 | 15 | N | T | *Mitochondrial complex I deficiency (MT-C1D)* | *252010* |
| *NUBPL* | 38 | 8 | L | P |  |  |
| *NUBPL* | 38 | 9 | D | Y |  |  |
| *NUBPL* | 38 | 13 | R | I |  |  |
| *OAT* | 35 | 1 | M | I | *Hyperornithinemia with gyrate atrophy of choroid and retina (HOGA)* | *258870* |
| *OAT* | 35 | 16 | R | H |  |  |
| *OAT* | 35 | 26 | A | V |  |  |
| *OAT* | 35 | 4 | G | E |  |  |
| *OAT* | 35 | 33 | Q | R |  |  |
| *OPA1* | 87 | 34 | S | N | *Optic atrophy 1 (OPA1)* | *165500* |
| *OPA1* | 87 | 32 | A | V |  |  |
| *OPA1* | 87 | 68 | A | V |  |  |
| *OPA1* | 87 | 69 | R | L |  |  |
| *OTC* | 32 | 1 | M | V | *Ornithine carbamoyltransferase deficiency (OTCD)* | 311250 |
| *OTC* | 32 | 26 | R | Q |  |  |
| *PANK2* | 46 | 1 | M | T | *Neurodegeneration with brain iron accumulation 1 (NBIA1)* | *234200* |
| *PANK2* | 46 | 35 | S | N |  |  |
| *PANK2* | 46 | 29 | T | A |  |  |
| *PANK2* | 46 | 10 | G | E |  |  |
| *PARS2* | 29 | 28 | R | S |  |  |
| *PCCA* | 52 | 42 | E | D | *Propionic acidemia type I (PA-1)* | 606054 |
| *PCK2* | 32 | 23 | S | X | *Mitochondrial phosphoenolpyruvate carboxykinase deficiency (M-PEPCKD)* | *261650* |
| *PDHA1* | 30 | 10 | R | P | *Pyruvate dehydrogenase E1-alpha deficiency (PDHAD)* | *312170* |
| *PDHX* | 53 | 23 | R | C | *Pyruvate dehydrogenase E3-binding protein deficiency (PDHXD)* | *245349* |
| *PDHX* | 53 | 5 | G | S |  |  |
| *PDHX* | 53 | 24 | R | G |  |  |
| *PDHX* | 53 | 41 | T | A |  |  |
| *PDHX* | 53 | 15 | R | H |  |  |
| *PMPCA* | 33 | 3 | R | Q | *Spinocerebellar ataxia, autosomal recessive, 2 (SCAR2)* | *213200* |
| *RARS2* | 16 | 12 | Q | R | *Pontocerebellar hypoplasia 6 (PCH6)* | *611523* |
| *RMND1* | 12 | 8 | T | M | *Combined oxidative phosphorylation deficiency 11 (COXPD11)* | *614922* |
| *RTN4IP1* | 40 | 3 | R | H | *Optic atrophy 10 with or without ataxia, mental retardation, and seizures (OPA10)* | *616732* |
| *RTN4IP1* | 40 | 23 | R | H |  |  |
| *SCO1* | 67 | 63 | P | L | *Mitochondrial complex IV deficiency (MT-C4D)* | 220110 |
| *SCO1* | 67 | 21 | G | S |  |  |
| *SCO2* | 41 | 36 | W | L | *Cardioencephalomyopathy, fatal infantile, due to cytochrome c oxidase deficiency 1 (CEMCOX1)* | *604377* |
| *SDHA* | 42 | 1 | M | L | *Mitochondrial complex II deficiency (MT-C2D)* | *252011* |
| *SDHB* | 28 | 3 | A | G | *Pheochromocytoma (PCC)* | *171300* |
| *SDHB* | 28 | 27 | R | X |  |  |
| *SDHD* | 56 | 50 | H | R | *Paragangliomas 1 (PGL1)* | 168000 |
| *SDHD* | 56 | 12 | G | S |  |  |
| *SDHD* | 56 | 1 | M | I |  |  |
| *SDHD* | 56 | 38 | R | G |  |  |
| *SDHD* | 56 | 42 | P | L |  |  |
| *SDHD* | 56 | 53 | D | Y |  |  |
| *SDHD* | 56 | 56 | L | P |  |  |
| *SDHD* | 56 | 30 | E | K |  |  |
| *SOD2* | 24 | 16 | V | A | *Microvascular complications of diabetes 6 (MVCD6)* | 612634 |
| *SUCLA2* | 52 | 39 | G | R | *Mitochondrial DNA depletion syndrome 5 (MTDPS5)* | *612073* |
| *SUCLA2* | 52 | 26 | R | Q |  |  |
| *TK2* | 33 | 24 | H | N | *Mitochondrial DNA depletion syndrome 2 (MTDPS2)* | 609560 |
| *TK2* | 33 | 11 | T | M |  |  |
| *TK2* | 33 | 22 | I | M |  |  |
| *TK2* | 33 | 4 | I | M |  |  |
| *TK2* | 33 | 27 | N | M |  |  |
| *TK2* | 33 | 9 | N | S |  |  |
| *TK2* | 33 | 33 | T | M |  |  |
| *TK2* | 33 | 15 | T | M |  |  |
| *TK2* | 33 | 17 | C | W |  |  |
| *TK2* | 33 | 15 | Y | N |  |  |
| *TK2* | 33 | 30 | S | G |  |  |
| *TK2* | 33 | 12 | S | G |  |  |
| *TRAP1* | 59 | 8 | R | S |  |  |
| *TTC19* | 70 | 3 | R | W | *Mitochondrial complex III deficiency, nuclear 2 (MC3DN2)* | *615157* |
| *TTC19* | 70 | 65 | W | X |  |  |
| *TWNK* | 31 | 3 | T | I | *Progressive external ophthalmoplegia with mitochondrial DNA deletions, autosomal dominant, 3 (PEOA3)* | *609286* |
| *TWNK* | 31 | 21 | A | P |  |  |
| *TWNK* | 31 | 2 | L | V |  |  |
| *TWNK* | 31 | 9 | R | G |  |  |
| *TXNRD2* | 36 | 36 | A | S | *Glucocorticoid deficiency 5 (GCCD5)* | *617825* |

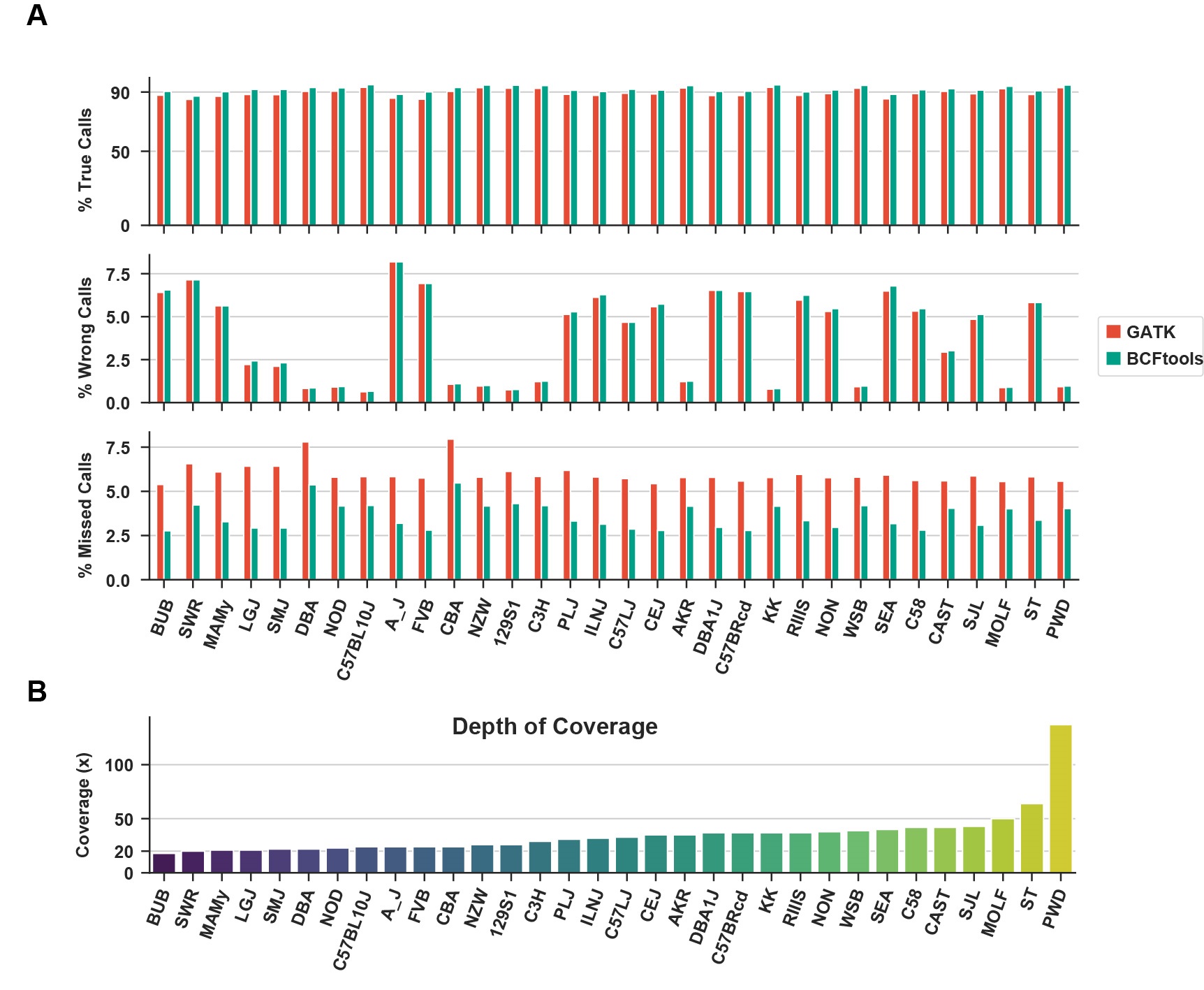

**Figure S1. The quality control of the called variants. (A)** A comparison of the results obtained using the BCFtools and GATK HaplotypeCaller pipelines. The MGI dbSNP database was used as the reference for the comparisons. For each of the 32 inbred strains, the percentage of correct, wrong or missed allelic calls is shown. (**B**) The depth of genomic sequence coverage (fold-genome) for the 32 inbred strains evaluated in (A).

frequency

|  | M | F |
| --- | --- | --- |
| 129S1 | 0 | 0 |
| AKR | 0 | 0 |
| A/J | 0 | 0 |
| B_C | 0 | 0 |
| BTBR | 0 | 0 |
| BUB | 0 | 0 |
| C3H | 0 | 0 |
| C57BRcd | 0 | 0 |
| C57BL10J | 0 | 0 |
| C57LJ | 0 | 0 |
| C57/6J | 0 | 0 |
| C58 | 0 | 0 |
| CBA | 0 | 0 |
| CEJ | 0 | 0 |
| DBA | 0 | 0 |
| FVB | 0 | 0 |
| ILNJ | 0 | 0 |
| KK | 0 | 0 |
| MAMy | 0 | 0 |
| NOD | 0 | 0 |
| NON | 0 | 0 |
| NZB | 0 | 0 |
| NZW | 0 | 0 |
| PLJ | 0 | 0 |
| SEA | 0 | 0 |
| SJL | 0 | 0 |
| SMJ | 0 | 0 |
| SWR | 0 | 0 |
| LPJ | 1 | 1 |

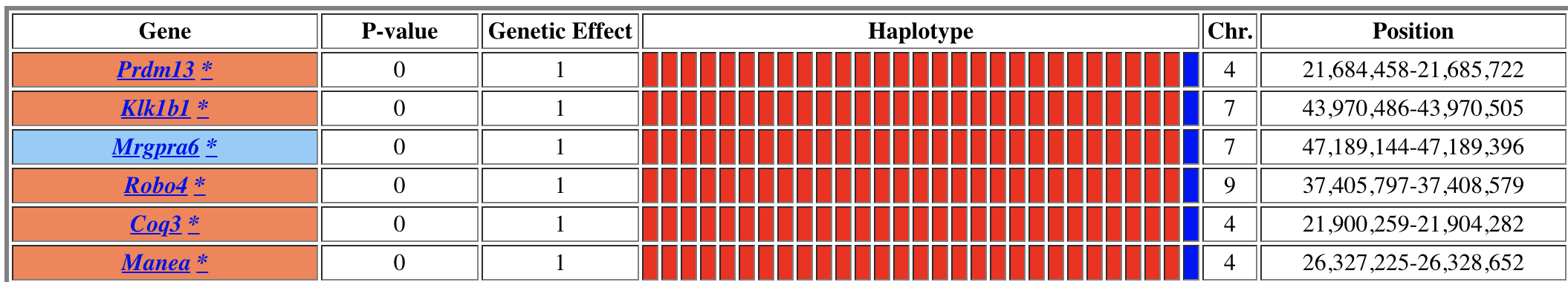

**Figure S2.** **Top:** The frequency of retinal hypopigmentation in 29 inbred mouse strains. The bar indicates that the percentage of male (blue) and female (red) LP/J mice with retinal hypopigmentation was 100%; while it was absent in the other 28 inbred strains evaluated. **Bottom:** HT-HBGCM output showing 6 genes with a perfectly correlated genetic pattern, which have a codon-changing SNP. The genes within the correlated haplotype blocks are indicated by their symbol; and a white, orange, or blue background indicates whether a SNP causes or does not cause an amino acid change; or if it affects a splice site, respectively. The haplotypic pattern is shown as colored rectangles that are arranged in the same order as the input data. Strains with the same colored rectangle have the same haplotypic pattern within the block. The p-values and genetic effect size were calculated as described [24].

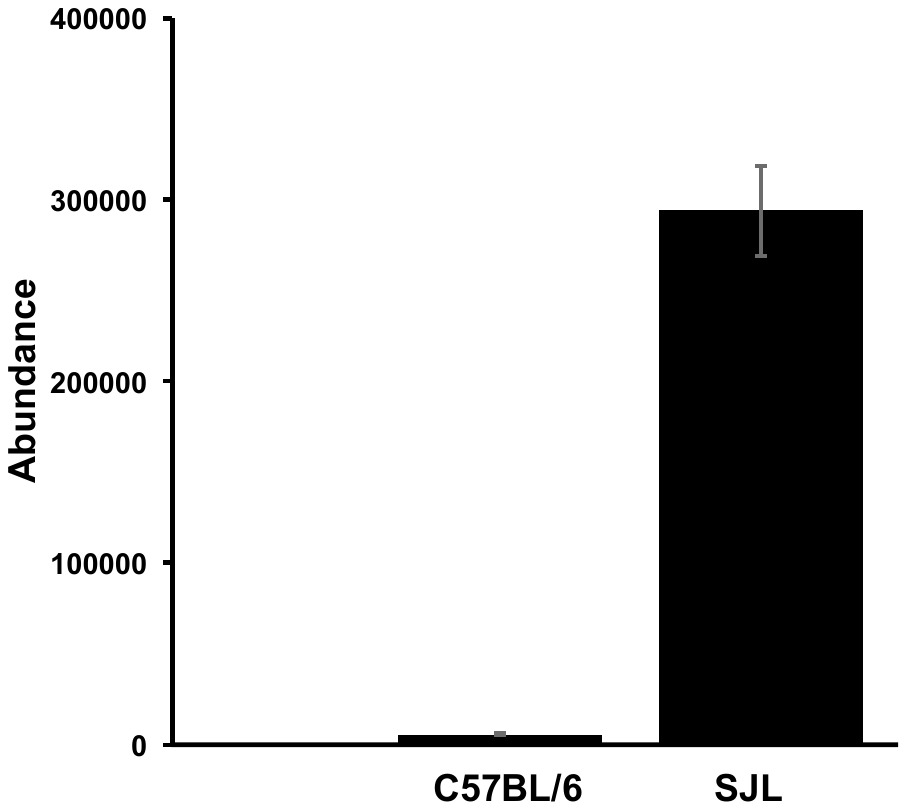

**Figure S3**. Succinylcarnitine levels were measured in livers obtained from C57BL/6 and SJL mice (n=5 per group) by LCMS analysis. Succinylcarnitine abundance in SJL liver was 50-fold greater than in C57BL/6 liver (p-value of 1.3 x 10^-10^).

|  | D508N | V271A | A266T | S247P | R110G | P88L | G23R |
| --- | --- | --- | --- | --- | --- | --- | --- |
| FVB | 0 | 1 | 1 | 1 | 1 | 1 | 0 |
| SJL | 0 | 1 | 1 | 1 | 1 | 1 | 0 |
| C57L/J | 0 | 0 | 0 | 0 | 0 | -1 | -1 |
| 129P2 | 0 | 0 | 0 | 0 | 0 | 0 | 0 |
| 129S1 | 0 | 0 | 0 | 0 | 0 | 0 | 0 |
| 129S5 | 0 | 0 | 0 | 0 | 0 | 0 | 0 |
| A/J | 0 | 0 | 0 | 0 | 0 | 0 | 0 |
| AKR | 0 | 0 | 0 | 0 | 0 | 0 | 0 |
| Balb/cJ | 0 | 0 | 0 | 0 | 0 | 0 | 0 |
| BTBR | 0 | 0 | 0 | 0 | 0 | 0 | 0 |
| C3H | 0 | 0 | 0 | 0 | 0 | 0 | 0 |
| C57BL/6J | 0 | 0 | 0 | 0 | 0 | 0 | 0 |
| C57BL/6NJ | 0 | 0 | 0 | 0 | 0 | 0 | 0 |
| C58 | 0 | 0 | 0 | 0 | 0 | 0 | 0 |
| CBA | 0 | 0 | 0 | 0 | 0 | 0 | 0 |
| CE/J | 0 | 0 | 0 | 0 | 0 | 0 | 0 |
| LP/J | 0 | 0 | 0 | 0 | 0 | 0 | 0 |
| NON | 0 | 0 | 0 | 0 | 0 | 0 | 0 |
| NU/J | 0 | 0 | 0 | 0 | 0 | 0 | 0 |
| NZB | 0 | 0 | 0 | 0 | 0 | 0 | 0 |
| PL/J | 0 | 0 | 0 | 0 | 0 | 0 | 0 |
| RF/J | 0 | 0 | 0 | 0 | 0 | 0 | 0 |
| RHJ | 0 | 0 | 0 | 0 | 0 | 0 | 0 |
| RIIIS | 0 | 0 | 0 | 0 | 0 | 0 | 0 |
| SM/J | 0 | 0 | 0 | 0 | -1 | 0 | 0 |
| C57BL/10J | 0 | 0 | 0 | 0 | 0 | 0 | 1 |
| C57BRcd | 0 | 0 | 0 | 0 | 0 | 0 | 1 |
| KK | 0 | 0 | 0 | 0 | 0 | 0 | 1 |
| MA/My | 0 | 0 | 0 | 0 | 0 | 0 | 1 |
| MRL | 1 | 0 | 0 | 0 | 0 | 0 | -1 |
| SWR | 1 | 0 | 0 | 0 | 0 | -1 | -1 |
| BUB | 1 | 0 | 0 | 0 | 0 | 0 | 0 |
| DBA/1J | 1 | 0 | 0 | 0 | 0 | 0 | 0 |
| DBA/2J | 1 | 0 | 0 | 0 | 0 | 0 | 0 |
| I/LnJ | 1 | 0 | 0 | 0 | 0 | 0 | 0 |
| LG/J | 1 | 0 | 0 | 0 | 0 | 0 | 0 |
| NOD | 1 | 0 | 0 | 0 | 0 | 0 | 0 |
| NZO | 1 | 0 | 0 | 0 | 0 | 0 | 0 |
| NZW | 1 | 0 | 0 | 0 | 0 | 0 | 0 |
| P/J | 1 | 0 | 0 | 0 | 0 | 0 | 0 |
| SEA | 1 | 0 | 0 | 0 | 0 | 0 | 0 |
| ST | 1 | 0 | 0 | 0 | 0 | 0 | 0 |

**Figure S4**. Codon changing SNPs in *Lactb*. The rows represent the indicated mouse strain, and the columns show the 5 codon changing SNPs. The alleles for each SNP are indicated by a colored box: blue for the reference (C57BL/6) strain, yellow for the alternative allele, and white is an unknown. FVB and SJ/L mice share a haplotype with unique SNP alleles at five sites: Arg110Gly and Pro88Leu are within the mitochondrial localization peptide; while *Val271Ala, Ala266Thr* and *Ser247Pro* SNps are within the sequence of the mature Lactb protein.

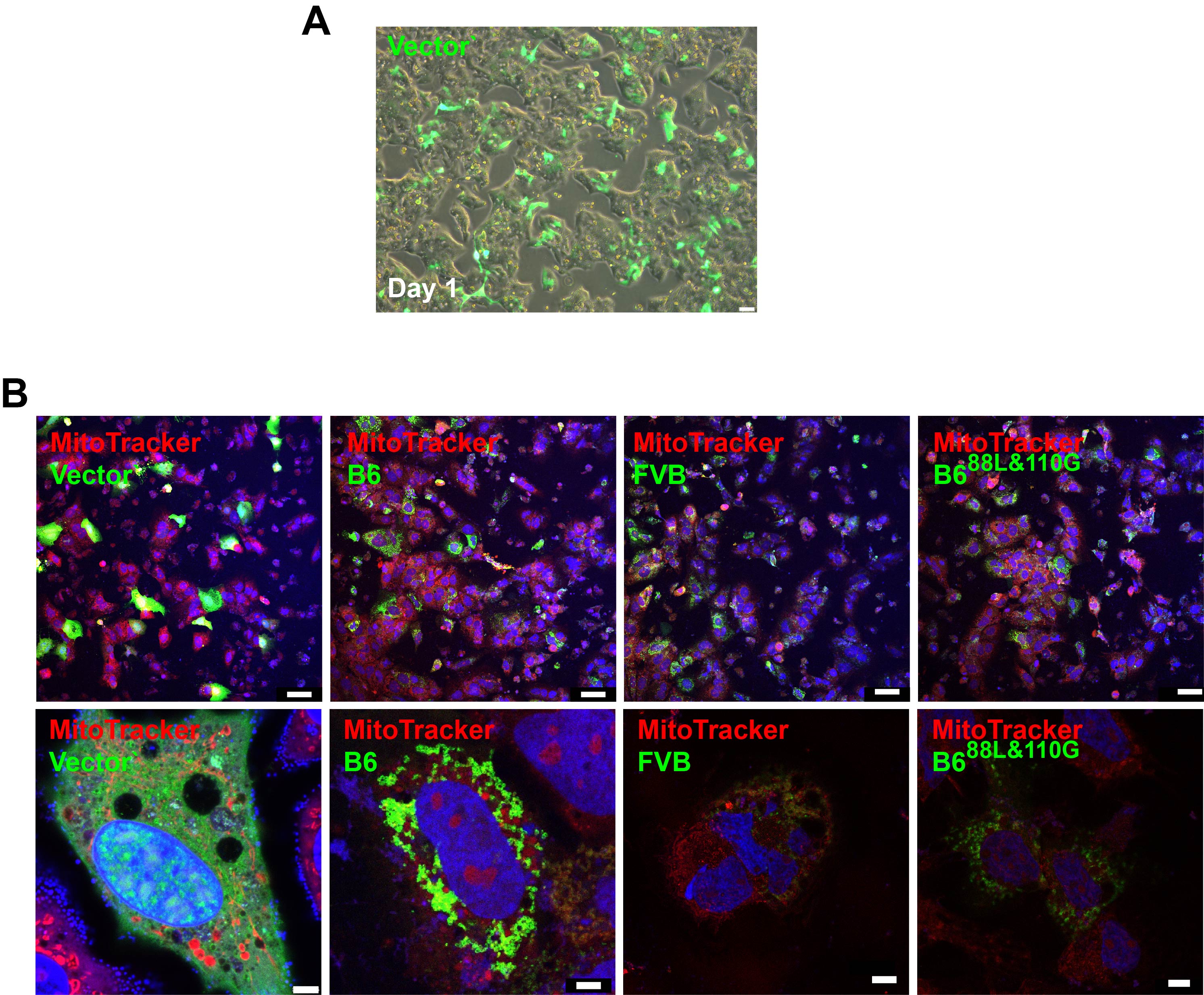

**Figure S5**. (**A**) A phase contrast image demonstrates that HepG2 cells were efficiently transfected with the control EGFP plasmid vector. (**B**) Images obtained 24 hours after HepG2 cells were transfected with the different plasmids (described in Figure 6A) indicate the different levels of expression and sub-cellular localization of the expressed fusion proteins. For each construct, the low magnification images are shown in the upper row (scale bar 50 um), and higher magnification images of single cells are shown in the lower row (scale bar 5 um). In these images, mitochondria are stained red, nuclei are stained blue, and the green color indicates the level of the LACTB-EGFP fusion protein expression. While the control EGFP-LACTB fusion protein is diffusely expressed in the cytoplasm, the C57BL/6 fusion protein has a more punctate pattern of expression that overlaps with mitochondria. The FVB EGFP-LACTB fusion protein has a lower level of expression; and the expression level of C57BL/6 fusion protein with FVB alleles at positions 88 and 110 resembles that of the FVB fusion protein.
